# Reactivating a schizophrenia risk gene-enriched prefrontal ensemble suppresses decision noise

**DOI:** 10.64898/2026.06.14.732113

**Authors:** Yusuke Iino, Hiroto Narita, Chika Shimizu, Shoi Shi

## Abstract

Schizophrenia genetics and single-cell transcriptomics implicate prefrontal excitatory neurons and synaptic programs, but how these vulnerabilities alter circuit dynamics, behavior, and pharmacological targetability remains unclear. We identify a clozapine-responsive prefrontal ensemble enriched for schizophrenia risk genes that stabilizes value-guided choice. NMDA receptor hypofunction increased value-independent decision noise and weakened the ensemble’s pre-choice transient, while clozapine rescued both. Chemogenetic inhibition of the ensemble increased decision noise, demonstrating causality. By integrating receptor affinities with mouse and human single-cell transcriptomes, we designed a rational multi-receptor antagonist cocktail that reactivated the ensemble and rescued decision noise without increasing NREM sleep time or NREM delta power, in contrast to clozapine. These findings link schizophrenia genetics to cortical ensemble dynamics and establish ensemble-targeted pharmacology as a strategy for ameliorating pathological computations.

**One-Sentence Summary:** Targeting a schizophrenia risk gene-enriched prefrontal ensemble suppresses decision noise without clozapine-like sedation.

## Main Text

Human genetics has reshaped the biology of schizophrenia. Common- and rare-variant studies (*1*, *2*), together with human single-cell transcriptomics (*3*), have converged on prefrontal cortical neurons, glutamatergic and synaptic programs, and cell-type-specific molecular vulnerability. Yet these molecular maps remain functionally incomplete. They identify genes, cell classes and molecular pathways associated with disease risk, but do not reveal how vulnerable cortical programs operate during behavior, which computations they support, or how they can be selectively recruited by pharmacological intervention. Bridging this gap requires decomposing psychiatric abnormalities into computationally defined behavioral phenotypes that can be quantified in humans (*4*, *5*), modeled across species, and causally tested in animal models.

Decision noise is one such computationally defined phenotype. Controlled stochasticity is essential for adaptive behavior: animals must occasionally depart from the currently favored option to explore alternatives, update latent values and adapt to changing environments (*6*, *7*). However, when stochastic choice becomes excessive and value-independent, it can impair goal-directed behavior (*8*). Computational psychiatry has identified increased decision noise as a quantitative behavioral abnormality relevant to psychosis and schizophrenia (*9*, *10*). The circuit mechanisms that actively constrain decision noise, particularly within prefrontal cortical ensembles, remain poorly understood. It is therefore unknown whether disease-relevant cortical cell programs merely mark genetic vulnerability or actively stabilize value-guided decisions.

Clozapine provides a pharmacological entry point into this problem. It is uniquely effective for treatment-resistant schizophrenia (*11*, *12*), yet its therapeutic action is not readily explained by strong blockade of a single receptor (*12*, *13*). Instead, clozapine’s broad receptor-binding profile raises the possibility that it recruits a distributed cortical ensemble shaped by combinatorial receptor expression. We reasoned that such a clozapine-responsive prefrontal ensemble could link schizophrenia-relevant synaptic cell programs to the stabilization of value-guided decision-making.

### NMDA receptor hypofunction increases value-independent stochastic choice

To quantify stochastic choice under changing reward contingencies, we trained mice in a two-choice probabilistic reversal-learning task (*14*). In each trial, mice initiated the trial by a center poke and then chose between two side ports associated with unequal reward probabilities. Reward contingencies reversed without explicit cues, requiring mice to use recent outcomes to update action values and adjust choices accordingly (Fig. 1A).

**Fig. 1.**
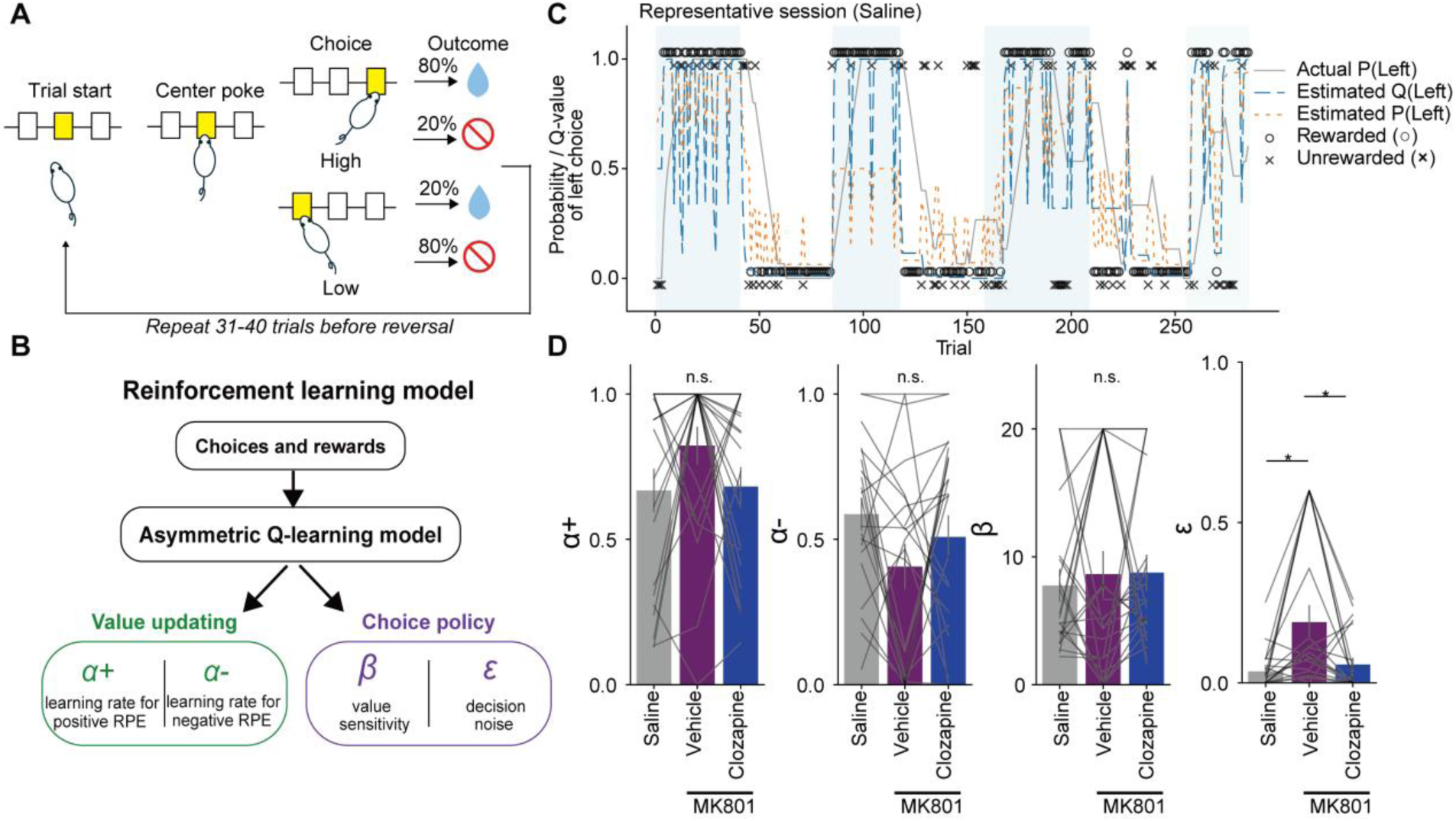
Clozapine rescues MK801-induced psychosis-relevant decision noise in a probabilistic reversal learning task. **(A)** Schematic of the reward-based decision-making task (two-alternative bandit task). Each trial was initiated by a center poke, after which mice chose between left and right reward ports. One option was associated with a high reward probability and the other with a low reward probability, and the reward contingencies reversed implicitly without explicit cues. **(B)** Reinforcement-learning model used to analyze behavior. Trial-by-trial choice behavior was fit with an asymmetric Q-learning model containing separate learning rates for positive and negative reward prediction errors (**α+** and **α-**), inverse temperature (**β**), and random choice parameter (**ε**). Choice probabilities were computed using a softmax decision rule with an additional random-choice component. **(C)** Representative behavioral sessions from saline-treated mice. Solid black lines indicate the actual reward probability assigned to the left option. Blue dashed lines indicate the estimated action value for the left option (Q(Left)), and orange dashed lines indicate the model-estimated probability of choosing the left option (P(Left)). Circles and crosses denote rewarded and unrewarded choices, respectively. **(D)** Fitted reinforcement-learning parameters across saline, MK801, and MK801 plus clozapine conditions. MK801 increased the value-independent random-choice parameter å, and this increase was rescued by clozapine. Bars indicate mean ± SEM, and gray lines connect paired measurements from individual mice (n = 21). Statistical comparisons were performed using one-way repeated-measures ANOVA followed by post hoc paired t tests with Holm correction. *P < 0.05; n.s., not significant.

We fit trial-by-trial choices with a reinforcement-learning model (*6*, *15*) that decomposed value updating and value use into positive and negative learning rates, inverse temperature, and a value-independent lapse or decision-noise parameter (Fig. 1B). Under vehicle conditions, mice gradually increased choices toward the higher-probability option after reversals, consistent with value-guided adaptation (Fig. 1C). Acute NMDA receptor blockade with MK801 increased apparently stochastic choices across blocks without detectably altering other parameters or behavioral measures (Fig. 1C, D and fig. S1). Model comparison supported the inclusion of a value-independent decision-noise parameter (fig. S2), and parameter-recovery analyses supported reliable estimation of this parameter under the behavioral conditions used here (figs. S3 and S4). To test whether increased dopaminergic tone produced the same computational signature, we examined dopamine transporter inhibition. This manipulation increased the learning rate for positive reward prediction errors and decreased the learning rate for negative reward prediction errors, while leaving inverse temperature and decision noise largely intact (fig. S5).

Thus, acute NMDA receptor hypofunction induces a selective, value-independent decision-noise phenotype, providing a computational readout for testing pharmacological rescue.

### Clozapine suppresses psychosis-relevant decision noise

We next asked whether clozapine, the most effective treatment for treatment-resistant schizophrenia (*11*, *12*), suppresses this computational abnormality. Clozapine improved value-guided choice in MK801-treated mice, restoring choice allocation toward the higher-probability option after reversals (Fig. 1D). Reinforcement-learning analysis showed that clozapine reduced MK801-induced decision noise toward vehicle levels (Fig. 1D). This rescue was dissociable from nonspecific effects on task performance. Clozapine-treated mice completed sufficient trials for model fitting, collected rewards reliably, and showed no improvement pattern consistent with a mere change in motivation or locomotor activity (fig. S1). By contrast, another antipsychotic, olanzapine, failed to rescue MK801-induced decision noise under matched task conditions (fig. S6). These results identify decision noise as a clozapine-sensitive computational phenotype in a mouse model of psychosis-relevant behavioral disruption.

### A clozapine-responsive prefrontal ensemble stabilizes value-guided choice

Because the medial prefrontal cortex is implicated in value-guided decision-making (*16*, *17*) and schizophrenia-related cognitive control (*18*, *19*), we asked whether clozapine engages a specific prefrontal neuronal ensemble during task performance. We combined clozapine treatment with activity-dependent labeling (*20*, *21*) to express inhibitory chemogenetic effector PSAM-GlyR (*22*) and Ca^2+^ indicator jGCaMP8f (*23*) in medial prefrontal cortex neurons recruited by clozapine (Fig. 2A, B).

**Fig. 2.**
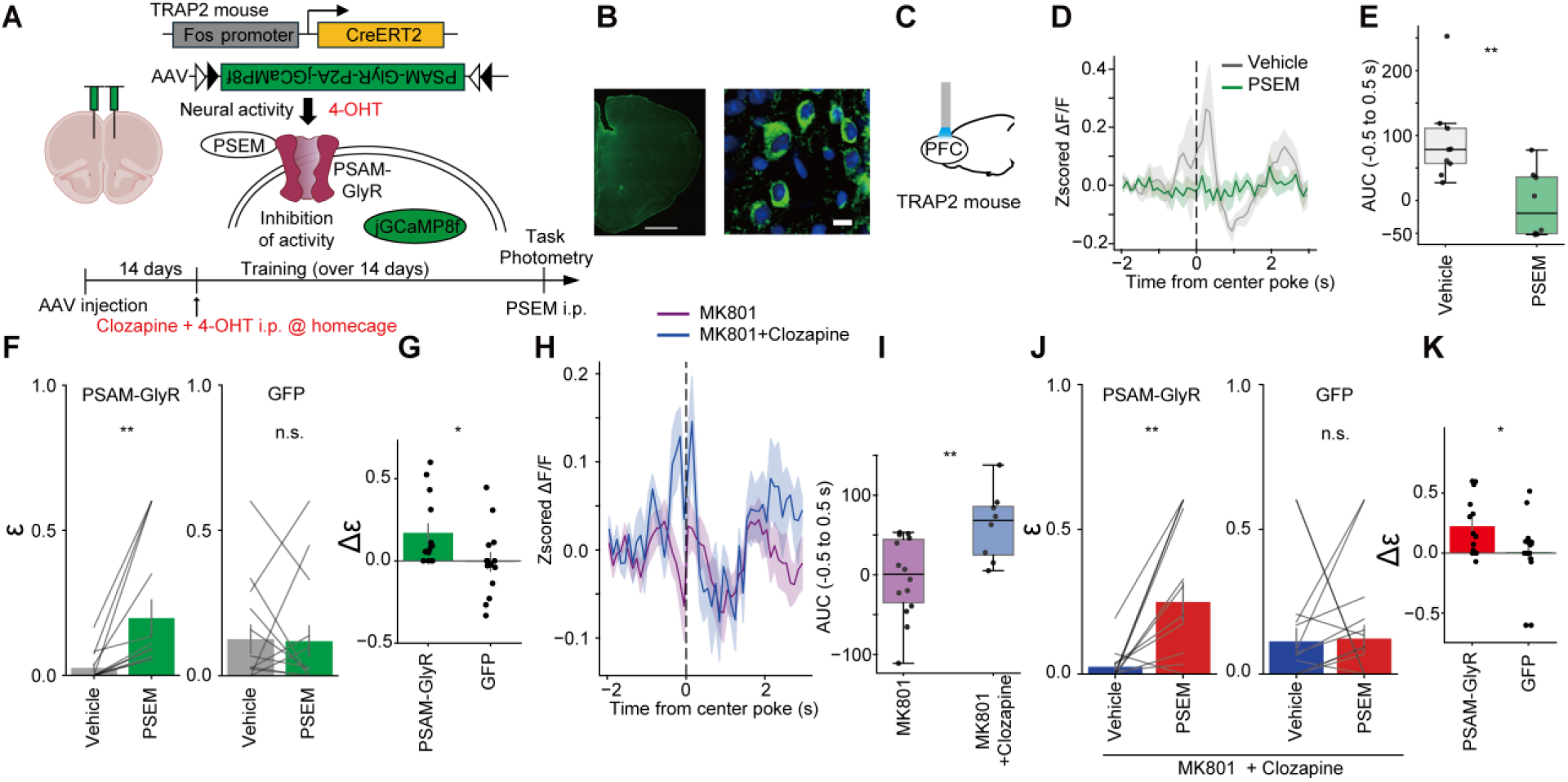
Clozapine-responsive prefrontal cells constrain psychosis-relevant decision noise. **(A)** Experimental strategy for activity-dependent targeting of clozapine-responsive PFC cells. In TRAP2 mice, clozapine plus 4-OHT was used to induce Cre-dependent expression of inhibitory PSAM-GlyR in clozapine-activated PFC cells. Subsequent administration of PSEM was used to inhibit these tagged cells during the two-armed bandit task. **(B)** Histological verification of viral expression and targeting in PFC. Scale bars, 1,000 µm (left) and 10 µm (right). **(C)** Schematic of fiber photometry recording from clozapine-responsive PFC cells expressing PSAM-GlyR together with jGCaMP8f. **(D, E)** Fiber photometry recording of clozapine-responsive PFC cells during bandit task with or without chemogenetic inhibition. Average z-scored calcium signals aligned to center poke are shown in **D**, and AUC (Area under curve) from −0.5 to 0.5 s around center poke is quantified in **E**. n = 9 mice (vehicle) and 8 mice (PSEM). **(F)** Inhibition of clozapine-responsive PFC cells during bandit task under baseline conditions in PSAM-GlyR (n = 14 mice) or GFP controls (n = 14 mice). **(G)** Within-subject change in å induced by PSEM relative to vehicle between PSAM-GlyR and GFP control mice. **(H, I)** Fiber photometry recording of clozapine-responsive PFC cells during the bandit task with or without clozapine under MK801 treatment. Average z-scored calcium signals aligned to center pokes are shown in **H**, and AUC is quantified in **I**. n = 14 mice (MK801) and 8 mice (MK801 + Clozapine). **(J)** Inhibition of clozapine-responsive PFC cells during bandit task under MK801 plus clozapine treatment in PSAM-GlyR (n = 14 mice) or GFP controls (n = 18 mice). **(K)** Within-subject change in å between PSEM and baseline conditions under MK801 plus clozapine treatment, comparing PSAM-GlyR and GFP control mice. Bars indicate group means, gray lines indicate paired individual data, and shaded areas indicate SEM. Within-mouse comparisons were performed using two-sided paired t tests in (**F** and **J**). Between-group comparisons were performed using two-sided Welch’s t tests in (**E**, **G**, **I** and **K**). *P < 0.05; **P < 0.01; n.s., not significant.

We then examined the activity dynamics of this ensemble using fiber photometry during task (Fig. 2C). Under control conditions, clozapine-responsive neurons showed a transient increase in activity during the trial-initiation period preceding side-port choice. These transients were abolished by PSEM administration (Fig. 2D, E), confirming effective chemogenetic suppression.

To test whether this ensemble contributes to choice stabilization, we chemogenetically inhibited clozapine-responsive neurons during baseline task performance. Inhibition of this ensemble increased decision noise even in the absence of MK801 (Fig. 2F, G). The effect was not accompanied by broad deficits in trial initiation, reward collection, or motor execution (fig. S7). These results indicate that clozapine-responsive medial prefrontal neurons are not merely markers of drug exposure but functionally constrain stochastic choice during normal value-guided behavior.

We next examined how this ensemble dynamic was altered under the MK801 and clozapine conditions. MK801 disrupted the pre-choice ensemble transient, whereas clozapine restored it (Fig. 2H, I). Then we asked whether this ensemble is required for the behavioral rescue by clozapine under MK801. Clozapine reduced the MK801-induced increase in decision noise, but this rescue effect was abolished when clozapine-responsive neurons were chemogenetically inhibited. By contrast, PSEM administration in GFP-expressing mice did not attenuate the clozapine-mediated rescue (Fig. 2J, K and fig. S8). These data suggest that clozapine suppresses decision noise by restoring transient activity of the prefrontal ensemble that precedes choice commitment.

### A single-cell GPCR model predicts that clozapine polypharmacology converges on schizophrenia-risk-enriched excitatory neurons

To convert clozapine’s broad receptor-binding profile into cell-type-resolved prediction of drug engagement, we constructed a single-cell GPCR signaling model that integrates drug–receptor affinity profiles (*24*) with receptor expression in single-cell transcriptomic data (*25*) (Fig. 3A–D). Because many antipsychotic drug targets signal through Gs- or Gi-coupled receptors, we estimated the net cAMP response of each cell to each antipsychotic drug. This model revealed heterogeneous predicted responses across prefrontal cortical cell clusters, with clozapine producing a response profile distinct from several other antipsychotics (Fig. 3E, F). In contrast, a parallel Gq-based calcium-response model did not capture the same clozapine-selective pattern (fig. S9). Thus, cAMP-linked Gs/Gi signaling provides a computational axis for identifying cells preferentially engaged by clozapine.

**Fig. 3.**
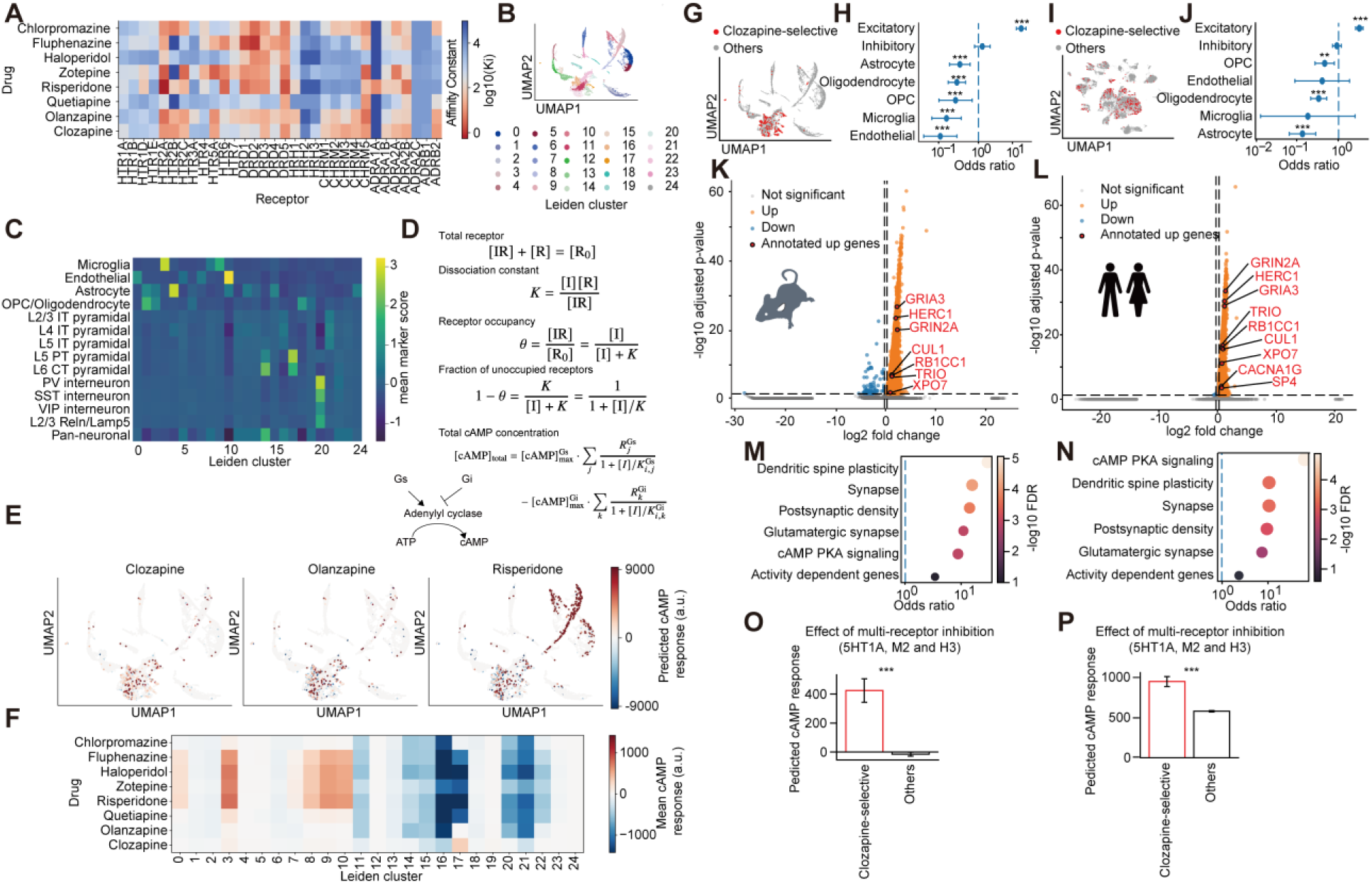
A single-cell cAMP model identifies schizophrenia-risk-gene-enriched clozapine-selective prefrontal neurons and a rational receptor-inhibition strategy. **(A)** Receptor-binding affinity matrix of antipsychotic drugs used for single-cell signaling prediction. Rows indicate antipsychotic drugs, columns indicate receptor targets, and color indicates −log10 Ki. **(B)** UMAP embedding of mouse mPFC single-cell RNA-seq data, colored by Leiden cluster. **(C)** Heatmap showing mean marker-gene scores across Leiden clusters, used to assign major neuronal and non-neuronal cell identities. **(D)** Schematic and equations for the receptor-occupancy-based cAMP model. Drug-induced cAMP responses were predicted from receptor affinity and single-cell receptor expression, with Gs-coupled receptors increasing and Gi-coupled receptors decreasing cAMP signaling. **(E)** UMAP plots showing predicted cAMP responses to clozapine, olanzapine, and risperidone in mouse mPFC cells. **(F)** Heatmap summarizing mean predicted cAMP responses for each antipsychotic drug across Leiden clusters. **(G)** UMAP visualization of predicted clozapine-selective cells in mouse mPFC. **(H)** Cell-type enrichment analysis of predicted mouse clozapine-selective cells. Odds ratios with confidence intervals are shown. **(I)** UMAP visualization of predicted clozapine-selective cells in human PFC. **(J)** Cell-type enrichment analysis of predicted human clozapine-selective cells. **(K, L)** Differential gene-expression analysis comparing predicted clozapine-selective cells with other cells in mouse mPFC (**K**) and human PFC (**L**). Volcano plots show significantly upregulated and downregulated genes, with annotated schizophrenia risk genes highlighted among upregulated genes. **(M, N)** Curated gene-set enrichment analysis of genes upregulated in predicted clozapine-selective cells in mouse (**M**) and human (**N**). Dot position indicates odds ratio, and color indicates −log10 FDR. **(O, P)** Predicted effects of simultaneous inhibition of 5HT1A, M2, and H3 receptors on cAMP responses in predicted clozapine-selective cells and other cells in mouse (**O**) and human (**P**) datasets. The multi-receptor inhibition pattern preferentially increased predicted cAMP responses in clozapine-selective cells. Cell-type enrichment in **H** and **J** was tested using two-sided Fisher’s exact tests with Benjamini–Hochberg FDR correction. Differential expression in **K** and **L** was tested using Wilcoxon rank-sum tests with Benjamini–Hochberg FDR correction. Curated gene-set enrichment in **M** and **N** was tested using two-sided Fisher’s exact tests with Benjamini–Hochberg FDR correction. Predicted cAMP responses in **O** and **P** were compared using two-sided Welch’s t tests. **P < 0.01, ***P < 0.001.

Using this model, we next predicted clozapine-selective cells, which are preferentially activated by clozapine, in mouse medial prefrontal cortex. These cells were significantly enriched in excitatory neuronal populations (Fig. 3G, H). We then applied the same modeling framework to human prefrontal cortex single-nucleus RNA-seq data (*26*). Despite species differences and dataset-specific technical variability, predicted clozapine-selective cells in the human cortex were again preferentially enriched among excitatory neurons (Fig. 3I, J). Thus, the model converged across mouse and human datasets on a conserved cellular target: a subset of prefrontal excitatory neurons.

We next asked whether these predicted clozapine-selective excitatory neurons carry molecular features relevant to schizophrenia pathophysiology (*3*). Differential expression analysis revealed that genes upregulated in predicted clozapine-selective cells were enriched for high-confidence schizophrenia risk genes (*2*) in both mice and human dataset (Fig. 3K, L). Notably, most of these differences remained significant in analysis within excitatory neurons (fig. S10). These findings suggest that the computationally defined clozapine-selective population is not only pharmacologically distinctive, but also aligned with the genetic architecture of schizophrenia.

Consistent with this interpretation, gene-set enrichment analysis showed that predicted clozapine-selective cells were enriched for synaptic and dendritic spine-related programs in both mice and human prefrontal cortex. Enriched terms included dendritic spine plasticity, synapse, postsynaptic density, glutamatergic synapse, and cAMP/PKA signaling, and activity-dependent genes (Fig. 3M, N). Together, these results define predicted clozapine-selective cells as a cross-species excitatory neuronal population enriched for schizophrenia risk genes and synaptic plasticity programs.

Finally, we used the model to search for receptor-inhibition combinations that could selectively recruit this schizophrenia-risk-gene-enriched clozapine-selective population without reproducing the full broad pharmacological profile of clozapine. We simulated multi-receptor inhibition patterns and ranked combinations by their ability to increase predicted cAMP signaling in clozapine-selective cells relative to other cells (fig. S11). The top-ranked combination included β2-adrenergic, M2 muscarinic, and H3 histamine receptor inhibition. Because systemic β2-adrenergic blockade has substantial translational liabilities (*27*), we prioritized the next-ranked combination, 5-HT1A, M2 muscarinic, and H3 histamine receptor inhibition, for experimental testing. Simulated 5-HT1A/M2/H3 inhibition preferentially increased predicted cAMP responses in clozapine-selective cells in both mouse and human prefrontal cortex data (Fig. 3O, P). The mRNAs of these receptors were preferentially expressed in clozapine-selective cells in both mice and human (fig. S12). These results provide a computational rationale for a clinically more tractable multi-receptor inhibition strategy to selectively recruit a conserved, schizophrenia-risk-gene-enriched population of clozapine-selective excitatory neurons.

### Rational receptor inhibition reactivates the clozapine-responsive ensemble and suppresses decision noise

We next tested whether the rational inhibitor cocktail identified by the GPCR signaling model recruits the clozapine-responsive prefrontal ensemble *in vivo*. Clozapine-responsive neurons were labeled in TRAP2 × Ai14 mice by pairing clozapine with 4-OHT, and mice were challenged two weeks later with clozapine, the inhibitor cocktail, or olanzapine (Fig. 4A). Clozapine rechallenge strongly reactivated tdTomato-positive neurons, as assessed by c-Fos immunostaining. The inhibitor cocktail induced a comparable overlap with the clozapine-tagged ensemble, whereas olanzapine showed significantly lower overlap (Fig. 4B, C). Thus, the rational inhibitor cocktail, but not olanzapine, preferentially reactivated the clozapine-responsive ensemble.

**Fig. 4.**
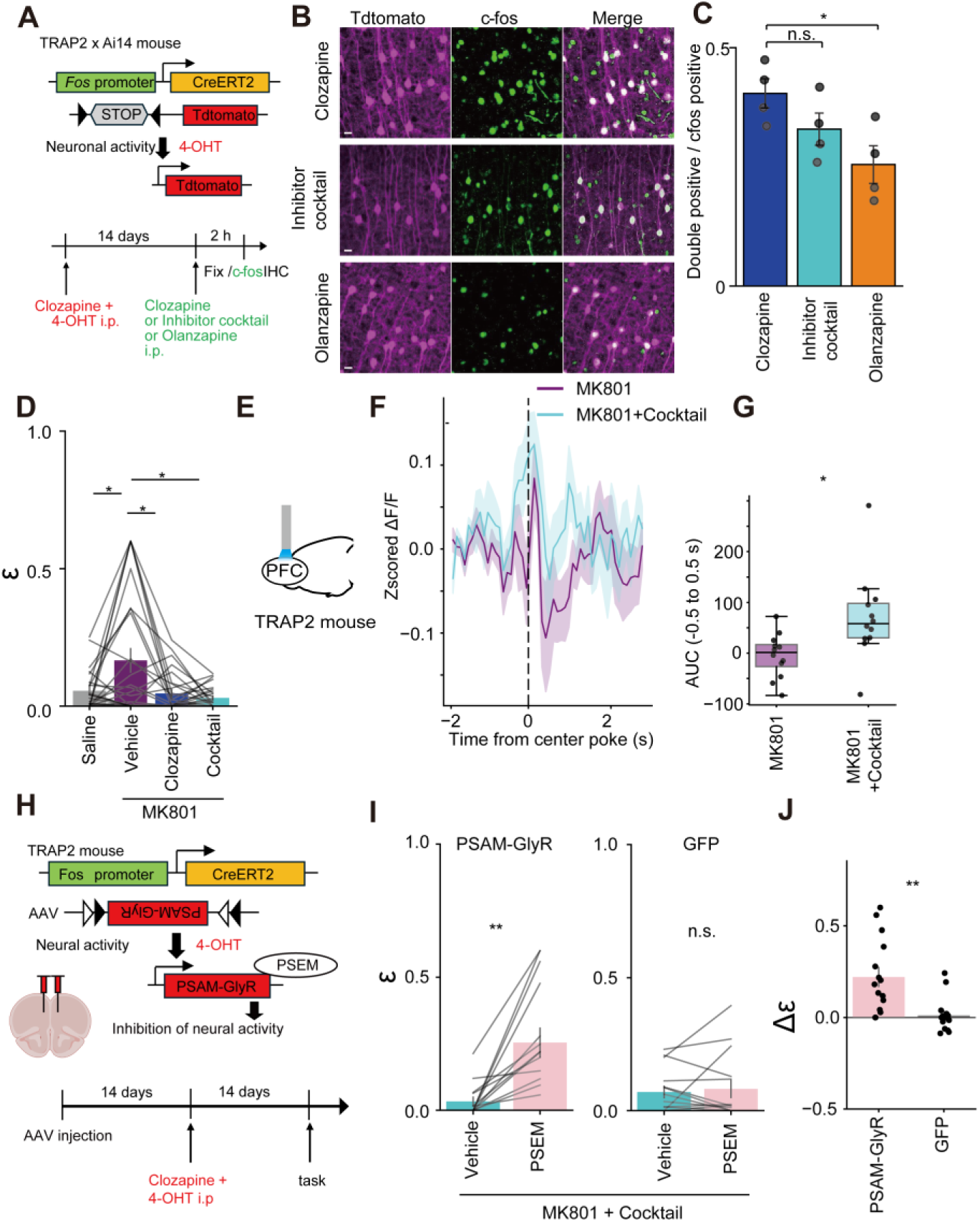
Rational pharmacological reactivation of clozapine-responsive prefrontal cells suppresses decision noise. **(A)** Experimental strategy for testing whether the rational inhibitor cocktail reactivates clozapine-responsive prefrontal cells. TRAP2 × Ai14 mice were treated with clozapine and 4-hydroxytamoxifen to permanently label clozapine-activated cells with tdTomato. After a washout period, mice received clozapine, the inhibitor cocktail, or olanzapine, and brains were fixed for c-Fos immunohistochemistry. **(B)** Representative images of tdTomato-labeled clozapine-responsive cells, c-Fos-positive cells, and merged images after clozapine, inhibitor cocktail, or olanzapine treatment. **(C)** Quantification of the fraction of c-Fos-positive cells that overlapped with tdTomato-labeled clozapine-responsive cells. n = 4 mice per condition. **(D)** Reinforcement-learning analysis of behavior in the two-alternative bandit task. MK801 increased the random choice parameter **ε**, and both clozapine and the inhibitor cocktail reduced this increase. Gray lines indicate individual mice (n = 26 mice). **(E-G)** Fiber photometry recording of clozapine-responsive prefrontal cells during the bandit task under MK801 and MK801 + inhibitor cocktail conditions. Schematic of fiber photometry recording is shown in (**E**). Average z-scored calcium signals aligned to center poke are shown in (**F**), and the area under the curve (AUC) from −0.5 to 0.5 s around center poke is quantified in (**G**). n = 12 mice per condition. **(H)** Experimental strategy for testing whether clozapine-responsive cells are required for the behavioral effect of the inhibitor cocktail. Cre-dependent PSAM-GlyR was expressed in clozapine-tagged mPFC cells of TRAP2 mice, allowing PSEM-mediated inhibition of this ensemble during the task. **(I)** Chemogenetic inhibition of clozapine-responsive PFC cells during the bandit task under MK801 + inhibitor cocktail conditions, comparing PSAM-GlyR mice (n = 15) and GFP control mice (n = 15). **(J)** Comparison of PSEM-induced changes in **ε** between PSAM-GlyR and GFP mice. Statistical comparisons were performed using one-way ANOVA followed by post hoc tests restricted to comparisons against the clozapine group with Holm correction for the two comparisons in (**C**), one-way repeated-measures ANOVA followed by post hoc paired t tests with Holm correction in (**D**), a two-sided Welch’s t test in (**G**) and (**J**), and two-sided paired t tests in (**I**). *P < 0.05; **P < 0.01; n.s., not significant.

We then examined whether this ensemble recruitment was associated with rescue of the pre-choice transient and decision noise. In the probabilistic reversal-learning task, MK801 increased decision noise, whereas both clozapine and the inhibitor cocktail suppressed this increase (Fig. 4D). These effects were not accompanied by broad changes in task engagement or motor measures (fig. S13). Fiber photometry from clozapine-tagged neurons showed that the inhibitor cocktail restored the pre-choice activity transient that was reduced under MK801 (Fig. 4E to G). These results indicate that the inhibitor cocktail recapitulates both the behavioral and ensemble-dynamic effects of clozapine.

Finally, we tested whether the clozapine-responsive ensemble is necessary for the effect of the inhibitor cocktail. We expressed PSAM-GlyR in clozapine-responsive neurons using TRAP2 mice and inhibited these neurons with PSEM during the MK801 plus inhibitor cocktail condition (Fig. 4H). PSEM increased decision noise in PSAM-GlyR mice but not in GFP controls (Fig. 4I), and the PSEM-induced change in decision noise was significantly larger in PSAM-GlyR mice than in GFP controls (Fig. 4J). Thus, the clozapine-responsive prefrontal ensemble is required for the inhibitor cocktail to suppress MK801-induced decision noise.

Together, these results support a causal chain linking receptor-level pharmacology, prefrontal ensemble dynamics, and stabilization of choice.

### Ensemble-targeted pharmacology dissociates cognitive rescue from clozapine-like sedation

A major limitation of clozapine is its broad side-effect profile, including sedation (*28*). Clozapine also alters sleep architecture in humans (*29*) and sleep–wake EEG states in rodents (*30*). We therefore asked whether the three-receptor cocktail could preserve the decision-stabilizing effect of clozapine while avoiding clozapine-like sleep-state changes. EEG/EMG recordings showed that clozapine increased non-rapid eye movement sleep and enhanced delta power, consistent with a sedative brain-state shift (Fig. 5A-D and fig. S14A-D). In contrast, the inhibitor cocktail did not increase NREM sleep and instead reduced delta power (Fig. 5E-H and fig. S14E-H).

**Fig. 5.**
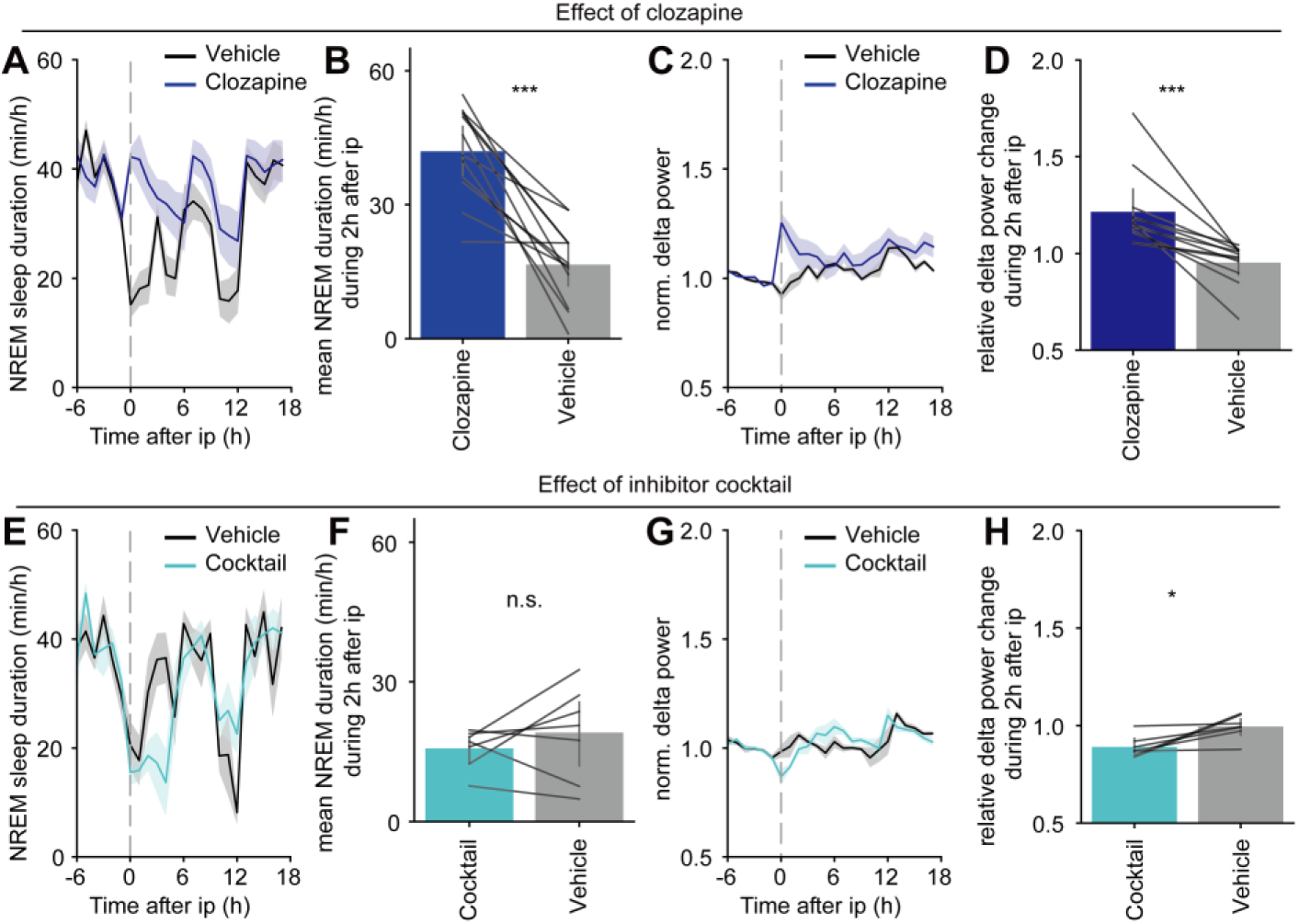
The rational inhibitor cocktail does not reproduce clozapine-induced NREM sleep and delta-power enhancement. **(A to D)** Effects of clozapine on NREM sleep and NREM delta power. **(A)** Time course of NREM sleep duration after intraperitoneal administration of vehicle or clozapine. **(B)** Quantification of mean NREM sleep duration during the first 2 h after injection. Gray lines indicate paired measurements from individual mice. **(C)** Time course of normalized NREM delta power after vehicle or clozapine administration. **(D)** Quantification of relative NREM delta power during the first 2 h after injection. n = 12 mice. **(E to H)** Effects of the rational inhibitor cocktail on NREM sleep and NREM delta power. **(E)** Time course of NREM sleep duration after intraperitoneal administration of vehicle or the inhibitor cocktail. **(F)** Quantification of mean NREM sleep duration during the first 2 h after injection. **(G)** Time course of normalized NREM delta power after vehicle or inhibitor cocktail administration. **(H)** Quantification of relative NREM delta power during the first 2 h after injection. n = 7 mice. Data are shown as mean ± SEM. Statistical analyses were performed using two-sided Wilcoxon signed-rank tests. **P < 0.01, ***P < 0.001; n.s., not significant.

Together, these findings indicate that clozapine’s suppression of psychosis-relevant decision noise can be separated from clozapine-like sedation. More broadly, they establish a framework for ensemble-targeted pharmacology, in which conserved receptor-expression logic is used to recruit a disease-relevant cortical ensemble and stabilize a computational dimension of behavior.

## Discussion

Human schizophrenia genetics and single-cell transcriptomics have converged on prefrontal excitatory neurons, glutamatergic signaling and synaptic programs as major components of disease biology (*1–3*). However, these molecular findings have remained difficult to connect to real-time cortical dynamics, behavior and pharmacological rescue. Here, we identify a clozapine-responsive medial prefrontal ensemble that stabilizes value-guided choice (Fig. 2) and is enriched for schizophrenia risk genes (Fig. 3G-N). NMDA receptor hypofunction increased value-independent decision noise (Fig. 1C, D) and weakened a pre-choice transient in this ensemble (Fig. 2H, I), whereas clozapine rescued both. Chemogenetic inhibition increased decision noise, demonstrating that this population is functionally required for decision stabilization (Fig. 2F, G). A rational multi-receptor antagonist cocktail reactivated the ensemble and rescued decision noise (Fig. 4) without reproducing clozapine-like sleep-state effects (Fig. 5). These findings link human schizophrenia genetics, cortical ensemble dynamics and pharmacological control of a computational phenotype.

Our results position decision noise as a bridge between computational psychiatry and circuit neuroscience. Controlled stochasticity supports exploration and adaptation, but excessive value-independent stochasticity can degrade the influence of learned values on action selection (*4*, *6–8*). By separating learning rates, inverse temperature and value-independent decision noise, our modeling suggests that NMDA receptor hypofunction prominently destabilizes choice rather than simply impairing value updating. This differs from dopamine-dependent mechanisms implicated in other psychosis-relevant computations including discrimination learning and perceptual inference (*31*, *32*). In our task, dopaminergic enhancement preferentially altered learning-rate parameters without producing a comparable increase in value-independent decision noise, suggesting that distinct psychosis-relevant perturbations can map onto separable computational dimensions. Although acute pharmacological manipulations in mice cannot be equated with clinical subtypes, this dissociation may provide a framework for interpreting part of the heterogeneity of schizophrenia-related decision-making abnormalities. Consistent with this interpretation, MK801 reduced a pre-choice activity transient in clozapine-responsive PFC cells, whereas clozapine restored this transient together with value-guided behavioral control. Moreover, inhibiting the same ensemble increased decision noise under baseline conditions and prevented clozapine-mediated rescue under NMDA receptor hypofunction (Fig. 2). These findings suggest that clozapine-responsive PFC cells provide a cortical stabilizing signal during action selection, allowing learned values to more reliably guide choice. Thus, clozapine’s rescue of unstable behavior may involve reinstatement of specific prefrontal ensemble dynamics rather than global enhancement of learning or nonspecific suppression of behavior.

In humans, schizophrenia is associated with abnormal activation of prefrontal cognitive-control networks during executive tasks, and reinforcement-learning fMRI studies have implicated prefrontal and anterior cingulate signals in altered reward-guided learning (*18*, *33*). Notably, clozapine, but not haloperidol, has been reported to restore task-activated anterior cingulate blood-flow patterns in schizophrenia (*34*). Thus, our finding that clozapine restores a pre-choice PFC ensemble transient in mice may provide a circuit-level entry point into prefrontal functional abnormalities observed in patients. This cross-species convergence does not imply a one-to-one homology between mouse medial PFC ensembles and human prefrontal networks, but it suggests that clozapine-sensitive prefrontal dynamics may represent an experimentally accessible circuit motif underlying impaired cognitive control in schizophrenia.

A major implication is that the clozapine-responsive ensemble is both pharmacologically defined and genetically relevant to schizophrenia. Human single-cell analyses supported conservation of the receptor-expression logic that defined the mouse clozapine-responsive population, and clozapine-responsive excitatory cortical cells were enriched for synaptic programs and schizophrenia risk genes (Fig. 3). This places clozapine-responsive prefrontal cells at the intersection of genetic liability, cortical cell identity and real-time decision dynamics. These findings also connect with the glutamate hypothesis of schizophrenia (*35*) and the clinical profile of clozapine in treatment-resistant schizophrenia (*11*). NMDA receptor hypofunction disrupted activity in this genetically vulnerable ensemble, while clozapine and the rational cocktail restored behavioral stability. Because treatment-resistant schizophrenia is not fully explained by canonical dopamine D2 receptor blockade (*12*), recruitment of a prefrontal synaptic ensemble may represent one mechanism through which clozapine stabilizes cognition. This does not imply that treatment resistance is a unitary glutamatergic disorder, but defines one experimentally tractable computation through which NMDA hypofunction, schizophrenia risk and clozapine-sensitive cortical mechanisms converge.

The rational cocktail supports an ensemble-centered view of pharmacology. Recent studies have shown that psychoactive drug actions can be decomposed into cell-type- or ensemble-specific components: Munguba et al. used cell-type transcriptomics to identify GPCR combinations that mimic antidepressant-relevant circuit mechanisms (*36*), whereas Muir et al. showed that reactivation of psychedelic-responsive prefrontal neurons recapitulates anxiolytic effects without reproducing hallucinogenic-like effects (*37*). Extending this principle to multi-receptor antipsychotic pharmacology, we integrated antipsychotic receptor-binding profiles with single-cell receptor expression to infer a clozapine-responsive ensemble and identify a receptor combination that selectively recruits it. This cocktail restored pre-choice ensemble dynamics and decision stability yet did not reproduce clozapine-like increase in NREM sleep or NREM delta power; instead, it reduced delta power. These findings suggest that stabilization of pathological decision noise can be dissociated, at least in part, from clozapine-like NREM sleep-state effects. More generally, receptor-transcriptome integration may enable the ensemble-level actions of multi-receptor drugs to be reverse-engineered and redesigned around specific circuit computations. In this framework, the relevant pharmacological unit is not an individual receptor, but a receptor-expression pattern that biases signaling within a functionally defined ensemble.

Several limitations remain. Acute MK801 treatment does not capture the full genetic, developmental and clinical heterogeneity of schizophrenia, and decision noise is only one dimension of psychosis-relevant behavior. Although human transcriptomic analyses support conservation and schizophrenia-risk-gene enrichment, they do not prove that the same functional ensemble exists in the human prefrontal cortex. Receptor-transcriptome modeling also cannot fully capture receptor protein abundance, ligand tone, pharmacokinetics or network context. Future studies using spatial transcriptomics, chronic genetic risk models and patient-derived systems will be needed to test the translational relevance of this ensemble more directly.

Together, our findings identify a schizophrenia-risk-gene-enriched prefrontal ensemble that is disrupted by NMDA receptor hypofunction and required for stabilization of decision noise. These results provide a mechanistic bridge from human schizophrenia genetics to cortical ensemble dynamics and establish ensemble-targeted pharmacology as a strategy for ameliorating pathological computations while minimizing broader brain-state side effects.

## Supporting information

Data S1. Mouse clozapine-selective cell DEG table

Data S2. Human clozapine-selective cell DEG table

Data S3. Mouse excitatory-cell DEG table

Data S4. Human excitatory-cell DEG table

## Acknowledgments

We express our gratitude to the staff of the Administrative Department and Animal Facility at IIIS. We thank all members of IIIS, especially M. Yanagisawa for insightful comments, and T. Sakurai, Y. Oishi and S. Honjo for their assistance in arranging experimental space. We also thank all members of the Shi Laboratory at IIIS, particularly K. Yoshida and M. Katori for valuable discussions on mathematical modeling, and M. Shiraishi and H. Tanaka for technical assistance.

## Author contributions

Conceptualization: Y.I.; Methodology: Y.I.; Investigation: H.N., C.S. and Y.I.; Software: Y.I.; Formal analysis: Y.I..; Resources: Y.I., and S.S.; Visualization: Y.I.; Funding acquisition: Y.I., and S.S.; Project administration: Y.I. and S.S.; Supervision: S.S.; Writing-original draft: Y.I.; Writing-review & editing: Y.I., and S.S.

## Data, code, and materials availability

All data are available in the main text or the supplementary materials. The code used for reinforcement learning model analysis, single-cell RNA-seq analysis, and intracellular signaling simulations will be available at https://github.com/yiino1222/RLmodel-and-snRNAseq-analysis and archived on Zenodo upon publication.

## Materials and Methods

All animal procedures followed the guidelines of the Institutional Animal Care and Use Committee of the University of Tsukuba.

### Single-cell RNA-seq preprocessing

Single-cell RNA-seq count matrices were analyzed using an AnnData/Scanpy-based workflow. For each dataset, gene symbols were converted to uppercase and duplicate gene names were made unique before downstream analysis. When multiple matrices were analyzed together, cells were concatenated across datasets using an outer join on genes, and dataset-level metadata, including dataset identifier, species, and brain region, were retained for downstream stratification. Cells with fewer than 200 detected genes or more than 6,000 detected genes were excluded, and genes detected in at least one cell were retained. Counts were normalized to 10,000 counts per cell and log-transformed. The log-normalized expression matrix was stored as the primary layer for differential-expression analyses. Where indicated, total counts were regressed out and expression values were scaled with a maximum value of 10. Highly variable genes were annotated using the top 4,000 genes but were not used to remove genes from receptor-focused or differential-expression analyses.

Dimensionality reduction was performed using principal component analysis (PCA; 50 components). A nearest-neighbor graph was constructed using 15 neighbors and 50 principal components, followed by UMAP embedding for visualization. Louvain and Leiden community-detection algorithms were applied for unsupervised clustering. GPU-accelerated routines were used when available; otherwise, the same preprocessing parameters and statistical definitions were applied using CPU-based Scanpy functions.

### Receptor binding affinity data

Receptor binding affinities (Ki values) for antipsychotic drugs were obtained from the Psychoactive Drug Screening Program (PDSP) Ki Database (*24*). For each drug–receptor pair, reported Ki values were converted to molar units and log-transformed where appropriate. When multiple Ki values were reported for the same receptor, the median value was used. Receptors with missing affinity data were excluded from modelling.

### Mouse scRNA-seq analysis and pharmacological response modeling

For mouse scRNA-seq data (*25*), receptor-expression features were extracted for a curated panel of antipsychotic-relevant G-protein-coupled receptors (GPCRs), including serotonergic, dopaminergic, histaminergic, muscarinic, adrenergic, and adenosine receptors. Raw receptor counts were retained in per-cell metadata fields when available. For pharmacological modeling, the receptor-expression matrix was normalized to 10,000 counts per cell. A drug-by-receptor affinity matrix was then used to calculate model-derived cAMP- and Ca-associated responses for each cell and each drug. These response values were used as computational phenotypes for identifying clozapine-responsive and clozapine-selective cells; they should be interpreted as receptor-expression- and affinity-weighted response scores rather than direct biochemical measurements

### cAMP-response model

The receptor panel was stratified by canonical G-protein coupling using the receptor-type annotation table. Let *E*_*i*,*r*_denote the normalized expression of receptor (r) in cell (i), (C) denote the modeled drug concentration, and *K*_*d*,*r*_ denote the affinity value for drug (d) and receptor (r) from the drug-by-receptor matrix. For Gs- and Gi-coupled receptors, the drug-adjusted residual receptor signal was calculated as

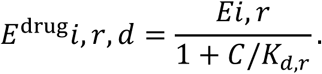

Basal cAMP-related signaling was defined as the difference between total Gs-coupled and total Gi-coupled receptor expression:

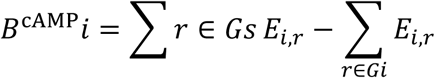

The modeled cAMP response for cell (i) and drug (d), stored as cAMP_<DRUG>, was calculated as the drug-adjusted Gs-minus-Gi signal minus the basal Gs-minus-Gi signal:

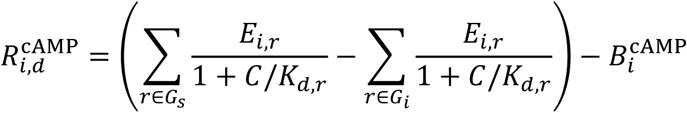

Thus, the cAMP score represents the modeled drug-induced change from each cell’s basal Gs-minus-Gi receptor-expression balance. Under this formulation, reduced contribution from Gi-coupled receptors shifts the score upward, whereas reduced contribution from Gs-coupled receptors shifts the score downward.

### Ca-response model

For Gq-coupled receptors, basal Ca-related signaling was defined as the total normalized expression of Gq-coupled receptors:

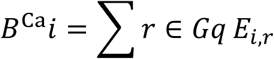

The drug-weighted Gq signal was calculated directly from the affinity matrix as ∑_*r*∈*Gq*_ *E*_*i*,*r*_ /*K*_*d*,*r*_. The modeled Ca response for cell (i) and drug (d), stored as Ca_<DRUG>, was therefore

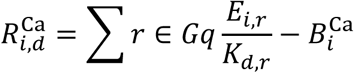

Ca and cAMP response scores were calculated for clozapine and for each comparator antipsychotic in the drug-by-receptor matrix, enabling cell-level comparison of clozapine-associated response patterns with those predicted for other antipsychotic drugs.

### Definition of clozapine-responsive and clozapine-selective cells

Clozapine-responsive cells were defined from the modeled cAMP response to clozapine. Cells with cAMP_CLOZAPINE > 10 were annotated as clozapine-activated, and cells with cAMP_CLOZAPINE < −10 were annotated as clozapine-inhibited. A separate clozapine-selective label was used to identify cells with a cAMP response preferentially associated with clozapine compared with other antipsychotic drugs.

For the selectivity calculation, all cAMP-response columns corresponding to non-clozapine drugs were first collected as comparator responses. For each cell, the mean comparator response was calculated as

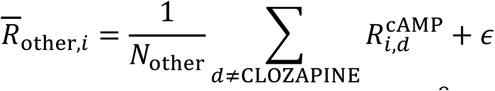

where *N*_other_ is the number of non-clozapine drugs and *ε* = 10^−9^ was added to avoid division by zero. Clozapine selectivity was then calculated as the squared ratio

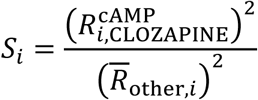

Cells were classified as clozapine-selective when (S_i) exceeded the prespecified selectivity threshold and the clozapine cAMP response was positive:

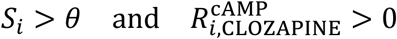

In the standardized pipeline configuration, the modeled drug concentration was set to 1,000 and the selectivity threshold *θ* was set to 1.5. This definition excludes cells with large-magnitude but negative clozapine cAMP responses from the clozapine-selective class. The resulting binary clozapine-selective annotation was used for downstream differential-expression analysis, enrichment analysis, and comparison with other antipsychotic-response profiles.

### Inhibitor-cocktail cAMP-pattern analysis

Candidate inhibitor-cocktail patterns were evaluated by specifying a binary inhibition pattern across receptor columns. Receptors included in the cocktail were assigned a low effective Ki value (Ki_inhibited = 0.01 by default), whereas receptors not included in the cocktail were assigned a high effective Ki value (Ki_not_inhibited = 10000 by default). Pattern keys corresponded to GPCR expression columns, such as HTR1A_raw, and receptors absent from the pattern were treated as not inhibited. For each receptor column (r), an effective (K_r) was defined from this binary pattern, and the cAMP response was recalculated using the same Gs/Gi framework:

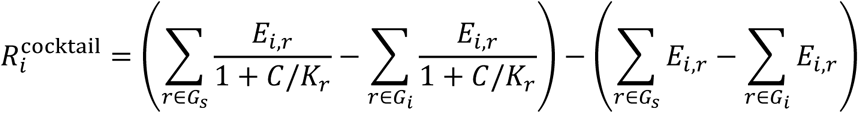

Only Gs- and Gi-coupled receptors present in the normalized GPCR expression matrix were included in this cocktail cAMP-pattern calculation. Cells were grouped by clozapine-selective status, and cAMP-response distributions were summarized by cell count, mean, median, and standard deviation. Differences between clozapine-selective and nonselective cells were evaluated using both Welch’s two-sample t-test and a two-sided Mann-Whitney U test.

### Human scRNA-seq analysis

Human scRNA-seq datasets (*26*) were processed using the same AnnData/Scanpy framework and pharmacological modeling definitions used for mouse data. Gene symbols were harmonized to uppercase, quality-control filters were applied, and expression matrices were normalized, log-transformed, embedded by PCA/UMAP, and clustered. Clozapine-selective response scores were inferred with the same receptor-affinity and receptor-coupling models after restricting calculations to receptors detected in the human expression matrix. When curated cell-type annotations were unavailable, broad cell classes were inferred from marker-gene module scores, including excitatory neurons, inhibitory neurons, astrocytes, oligodendrocyte-lineage cells, endothelial cells, and microglia. Annotated or inferred cell-type labels were used for enrichment analyses and for stratified interpretation of clozapine-selective populations.

### Differential-expression analysis

Differentially expressed genes (DEGs) between clozapine-selective and nonselective cells were identified using Scanpy’s rank_genes_groups function with the Wilcoxon rank-sum test applied to the log-normalized expression layer. The clozapine-selective annotation was encoded as a categorical grouping variable, and gene rankings, test statistics, log fold changes, adjusted p values, and detection fractions were extracted with scanpy.get.rank_genes_groups_df. Unless otherwise specified, genes were considered significant at a Benjamini-Hochberg false-discovery-rate (FDR) threshold of 0.05. An additional absolute log2 fold-change threshold of 0.25 was used for volcano-plot visualization and interpretation. For receptor-focused analyses, raw receptor-count columns were preferred when present; otherwise, expression values were extracted from the AnnData expression matrix. Receptor and marker expression differences between clozapine-selective and nonselective cells were tested using two-sided Mann-Whitney U tests, followed by Benjamini-Hochberg correction across tested genes.

### Cell-type and cluster enrichment analysis

Enrichment of clozapine-selective cells within annotated cell types, inferred cell types, or Leiden clusters was assessed using two-by-two contingency tables. Cell-type labels were obtained from dataset-provided annotations when available. For datasets in which annotations were unavailable, incomplete, or required validation, broad cell classes were inferred from marker-gene module scores calculated from the log-normalized expression matrix, together with visual inspection of marker expression on the UMAP embedding. Gene symbols were matched case-insensitively after conversion to uppercase, and mouse gene symbols were interpreted using uppercase ortholog-style names to maintain consistency with human datasets.

Broad cell classes were defined using established marker genes. Excitatory neurons were identified using glutamatergic and cortical projection-neuron markers, including SLC17A7, SLC17A6, SATB2, TBR1, CUX1, CUX2, RORB, THEMIS, BCL11B/CTIP2, FEZF2, FOXP2, TLE4, and PCP4. Inhibitory neurons were identified using GABAergic and interneuron-subclass markers, including GAD1, GAD2, SLC6A1, DLX1, DLX2, PVALB, SST, VIP, LAMP5, and RELN. Astrocytes were identified using AQP4, ALDH1L1, GFAP, SLC1A2, and SLC1A3. Oligodendrocyte-lineage cells were identified using oligodendrocyte markers, including MBP, MOG, PLP1, and MOBP, and oligodendrocyte precursor-cell markers, including PDGFRA and CSPG4. Microglia were identified using P2RY12, CX3CR1, AIF1, C1QA, and C1QB. Endothelial cells were identified using PECAM1, CLDN5, VWF, and FLT1. Pericytes and vascular smooth muscle cells were identified using PDGFRB, RGS5, and ACTA2. Ependymal or choroid-like cells were identified using FOXJ1 and TTR.

For each cell-type category or cluster, the contingency table compared the number of clozapine-selective and nonselective cells within that category with the corresponding numbers outside the category. Enrichment P values were calculated using two-sided Fisher’s exact tests. Odds ratios and 95% confidence intervals were estimated after applying a Haldane–Anscombe correction, in which 0.5 was added to each cell of the contingency table before calculating the log odds ratio and its standard error. P values across tested categories were adjusted using the Benjamini–Hochberg false-discovery-rate procedure. Enrichment results were visualized as forest plots on a log odds-ratio scale.

The same marker-based cell-identity framework was used for the excitatory-neuron-only DEG analysis described below.

### Excitatory-neuron-only DEG analysis

To determine whether clozapine-selective transcriptional differences were present within excitatory neurons, a subset analysis was performed after restricting the AnnData object to cells annotated or inferred as excitatory neurons by the cell-type marker framework described above. Briefly, cells were retained when dataset-provided annotations identified them as glutamatergic, excitatory, pyramidal, or cortical projection neurons, or when marker-based module scoring supported excitatory-neuron identity. Cells with stronger marker evidence for inhibitory neuronal, glial, vascular, or ependymal/choroid-like lineages were excluded using the same comparator marker sets used for cell-type enrichment analysis. Within the final excitatory-only subset, DEGs were recalculated between clozapine-selective and nonselective cells using the same Wilcoxon rank-sum framework, log-normalized expression layer, and multiple-testing correction described above. Excitatory-only volcano plots were generated using an FDR threshold of 0.05 and an absolute log2 fold-change threshold of 0.25 unless otherwise indicated. Selected disease-risk or mechanistic genes were labeled only by exact gene-symbol matching to avoid partial-match annotation artifacts.

### Curated pathway enrichment analysis for clozapine-selective DEGs

Targeted enrichment analysis was performed using manually curated gene sets selected a priori to support mechanistic interpretation of differentially expressed genes in clozapine-selective cells. Gene symbols were converted to uppercase before matching. When mouse datasets were analyzed, mouse gene symbols were mapped to uppercase human ortholog-style symbols when necessary so that the same curated gene sets could be applied consistently across mouse and human analyses. The tested gene universe was defined as the set of genes retained in the DEG result table after expression filtering and gene-symbol harmonization, rather than the whole genome. Genes included in a curated set but absent from the tested universe were retained in the reported definition of the gene set but excluded from contingency-table counts.

Five curated gene sets were analyzed. The synapse and synaptic transmission gene set was used to assess whether clozapine-selective DEGs were enriched for neuronal synaptic biology and included genes related to presynaptic vesicles, release machinery, postsynaptic density, adhesion, and glutamatergic synapses: SYN1, SYN2, SYN3, SYP, SNAP25, STX1A, STXBP1, VAMP2, SYT1, DLG4, SHANK1, SHANK2, SHANK3, HOMER1, NRXN1, NRXN2, NRXN3, NLGN1, NLGN2, NLGN3, GRIN1, GRIN2A, GRIN2B, GRIA1, GRIA2, CAMK2A, and CAMK2B. The cAMP/PKA signaling gene set was used to evaluate transcriptional changes related to the receptor-derived cAMP model and included adenylyl cyclases, PKA catalytic and regulatory subunits, phosphodiesterases, cAMP-responsive transcriptional mediators, EPAC proteins, and anchoring proteins: ADCY1, ADCY2, ADCY3, ADCY5, ADCY6, ADCY7, ADCY8, ADCY9, PRKACA, PRKACB, PRKACG, PRKAR1A, PRKAR1B, PRKAR2A, PRKAR2B, PDE1A, PDE1B, PDE1C, PDE2A, PDE3A, PDE3B, PDE4A, PDE4B, PDE4C, PDE4D, CREB1, ATF1, RAPGEF3, RAPGEF4, AKAP5, AKAP7, AKAP9, AKAP11, and AKAP12. The GPCR and monoamine receptor signaling gene set was used to compare DEG enrichment with the pharmacological receptor-response model and included antipsychotic-relevant receptors and proximal G-protein or arrestin signaling components: HTR1A, HTR2A, HTR2C, HTR6, HTR7, DRD1, DRD2, DRD3, DRD4, HRH1, HRH3, CHRM1, CHRM2, CHRM3, CHRM4, CHRM5, ADRA1A, ADRA2A, ADORA2A, GNAI1, GNAI2, GNAI3, GNAS, GNAQ, GNA11, ARRB1, and ARRB2. The calcium signaling and neuronal excitability gene set was used to assess Ca-associated signaling and excitability-related DEG patterns and included voltage-gated calcium channels, calmodulin/calcineurin components, and intracellular calcium-release genes: CACNA1A, CACNA1B, CACNA1C, CACNA1D, CACNA1E, CACNA1G, CACNA1H, CACNA1I, CAMK2A, CAMK2B, CALM1, CALM2, CALM3, PPP3CA, PPP3CB, PPP3CC, RYR1, RYR2, RYR3, ITPR1, ITPR2, and ITPR3. The immediate-early and activity-regulated transcription gene set was used to evaluate whether clozapine-selective cells showed an activated or activity-regulated expression signature and included activity-dependent transcriptional regulators and plasticity-associated genes: FOS, FOSB, JUN, JUNB, JUND, EGR1, EGR2, EGR3, EGR4, NPAS4, ARC, IER2, IER3, NR4A1, NR4A2, NR4A3, and BDNF.

For the curated over-representation analysis, significant DEGs were separated into upregulated and downregulated gene lists using the same FDR and absolute log2 fold-change thresholds used for DEG interpretation. For each DEG direction and each curated gene set, a two-by-two contingency table was constructed using the number of significant DEG genes inside the curated set, significant DEG genes outside the curated set, nonsignificant background genes inside the curated set, and nonsignificant background genes outside the curated set. Enrichment P values were calculated using two-sided Fisher’s exact tests. Odds ratios were reported with Haldane-Anscombe correction when zero counts were present. P values across curated gene sets and DEG directions were adjusted using the Benjamini-Hochberg false-discovery-rate procedure. Curated gene sets were considered significantly enriched at FDR < 0.05. Nominal P values, adjusted P values, odds ratios, overlap counts, effective term sizes, and overlapping gene symbols were reported for transparency.

### Statistical analysis

Unless otherwise specified, statistical tests were two-sided. Differential-expression testing used Wilcoxon rank-sum tests as implemented in Scanpy, with Benjamini-Hochberg adjustment for multiple comparisons. Receptor-level and marker-level comparisons between clozapine-selective and nonselective cells used Mann-Whitney U tests with Benjamini-Hochberg correction. Cell-type and cluster enrichment analyses used Fisher’s exact tests with Benjamini-Hochberg correction across categories. Inhibitor-cocktail cAMP-pattern comparisons were summarized descriptively and tested using Welch’s two-sample t-test and the Mann-Whitney U test. For visualization, adjusted P values equal to zero were replaced by the smallest positive adjusted P value in the corresponding result table or by a numerical floor to permit log-scale plotting. Data are reported as cell counts, means, medians, standard deviations, odds ratios with 95% confidence intervals, log2 fold changes, and FDR-adjusted P values as appropriate.

### In silico screening of receptor inhibition patterns

To identify receptor combinations capable of selectively modulating clozapine-responsive cells, we performed an in silico screen of receptor inhibition patterns using the single-cell GPCR-expression matrix and the Gs/Gi cAMP-response model. The candidate receptor set was restricted to Gs- and Gi-coupled receptors detected in the single-cell expression matrix, yielding 17 receptors in total, consisting of 8 Gs-coupled and 9 Gi-coupled receptors.

For each receptor inhibition pattern, receptors included in the pattern were modeled as inhibited, whereas receptors not included in the pattern were left unchanged. Inhibited receptors were assigned an effective (K_i) of 0.01, and non-inhibited receptors were assigned a large effective (K_i), such that their contribution was minimally altered. Drug concentration was set to (C=1). For each cell, the predicted cAMP response was calculated as the difference between the post-inhibition Gs–Gi balance and the basal Gs–Gi balance.

We first assessed how selectivity changed as the number of inhibited receptors increased. All possible receptor inhibition patterns containing (k = 1) to (6) inhibited receptors were enumerated across the 17 candidate Gs/Gi-coupled receptors. This yielded 17, 136, 680, 2,380, 6,188, and 12,376 combinations for (k = 1), (2), (3), (4), (5), and (6), respectively. In total, 21,777 receptor inhibition patterns were evaluated in the saturation analysis. For each pattern, a selectivity score was defined as the mean predicted cAMP response in clozapine-responsive cells minus the mean predicted cAMP response in non-responsive cells:

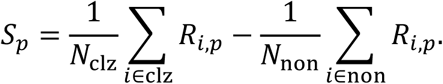

Where *R*_*i*,*p*_denotes the predicted cAMP response of cell (i) under inhibition pattern (p), and *N*_clz_and *N*_non_denote the numbers of clozapine-responsive and non-responsive cells, respectively. Positive values indicate preferential predicted cAMP modulation in clozapine-responsive cells. Saturation was evaluated by comparing the highest selectivity score and the mean selectivity score among top-ranked patterns across values of (k).

Based on this saturation analysis, we focused the final screen on three-receptor inhibition patterns. All possible three-receptor combinations among the 17 candidate receptors were enumerated, yielding 680 patterns. Each pattern was scored using the same selectivity index described above. Patterns were ranked by selectivity score, and receptor inclusion frequencies were calculated across top-ranked patterns to identify recurrent receptor motifs. Candidate receptor combinations were selected based on selectivity score, recurrence among top-ranked patterns, and pharmacological feasibility for in vivo testing.

### Animals

All animal experiments were conducted in accordance with institutional guidelines and approved by the Animal Care and Use Committee of the University of Tsukuba. Adult male C57BL/6J mice (8–12 weeks old) were used for behavioural and sleep experiments unless otherwise stated. TRAP2 (Fos-CreERT2) mice (*20*, *21*) were crossed with Ai14 reporter mice to generate TRAP2 × Ai14 animals for activity-dependent neuronal labelling. Mice were group-housed under a 12-h light/dark cycle (lights on at 5:00) with ad libitum access to food and water unless specified.

### Plasmid constructions and adeno-associated virus (AAV) preparations

The Cre-dependent AAV plasmid used for simultaneous chemogenetic inhibition and calcium imaging of TRAPed cells was custom synthesized by VectorBuilder. The construct was designed to express a 3×HA-tagged PSAM4-GlyR (*22*) and 6×His-tagged jGCaMP8f (*23*) from the EFS promoter in a Cre-dependent manner. The PSAM4-GlyR and jGCaMP8f coding sequences were separated by a P2A self-cleaving peptide to permit co-expression from a single transcript. The expression cassette was arranged in an inverted FLEXon configuration flanked by heterotypic lox sites, enabling Cre-mediated inversion and expression in recombined cells. The plasmid also contained a WPRE sequence to enhance transgene expression. The final sequence was synthesized and sequence-verified by VectorBuilder and used for AAV production. AAVs were produced and their titres were measured as described previously (*38*). In brief, plasmids for the AAV vector, pHelper (Stratagene) and pUCmini-iCAP-PHP.eB (#103005, Addgene) were transfected into HEK293 cells (AAV293, Stratagene). After three days, cells were harvested and AAVs were purified twice using iodixanol. The titers of AAVs were estimated using quantitative PCR.

### Surgery

Mice were anesthetized with isoflurane and placed in a stereotaxic frame. AAV was injected into the mPFC using a microsyringe and glass capillary. Approximately 500 nl of viral solution was injected per site at a rate of 100 nl/min at the following coordinates: AP, +1.98 mm relative to bregma; ML, ±0.30 mm relative to bregma; and DV, −1.80 mm from the dura. After viral infusion, the injection needle was left in place for 5 min before withdrawal to minimize backflow. For bilateral viral expression experiments, injections were performed in both hemispheres.

For fiber photometry experiments, an optical fiber cannula (R-FOC-BL400C-50NA, 5 mm; 400-µm core diameter, NA 0.5; RWD) was implanted unilaterally 200 µm above the viral injection site. The fiber tip was therefore positioned at DV −1.60 mm from the dura. The implanted hemisphere was counterbalanced across mice. The fiber cannula was secured to the skull with dental cement. Mice were allowed to recover after surgery before subsequent experiments.

### Drugs and pharmacological treatments

Clozapine (1 mg/kg; FUJIFILM Wako), olanzapine (1 mg/kg), MK801 (0.05 mg/kg; FUJIFILM Wako), GBR12909 (10 mg/kg; TCI), and uPSEM817 tartrate (0.6 mg/kg; Tocris) were used for pharmacological experiments. The receptor inhibitor cocktail consisted of WAY-100635 maleate (1 mg/kg), pitolisant (10 mg/kg), and (S)-(+)-dimethindene maleate (0.1 mg/kg), targeting 5-HT1A, histamine H3, and muscarinic M2 receptors, respectively.

Unless otherwise stated, drugs were dissolved in sterile saline. GBR12909 was dissolved in 10% DMSO in saline. For the receptor inhibitor cocktail, the three antagonists were mixed immediately before administration and injected as a single intraperitoneal injection. All drugs were administered intraperitoneally.

### TRAP2 labelling and viral expression

For activity-dependent labeling of clozapine-responsive neurons, TRAP2 × Ai14 mice received intraperitoneal injections of clozapine (1 mg/kg) and 4-hydroxytamoxifen (4-OHT; 50 mg/kg) in the home cage. 4-OHT was first dissolved in ethanol at 20 mg/ml by incubation at 37°C for 15 min. The 20 mg/ml 4-OHT solution was then mixed with corn oil at a 1:2 ratio and incubated at 37°C for 1 h to evaporate ethanol, yielding a final 4-OHT concentration of 10 mg/ml in corn oil. This procedure induced Cre-dependent tdTomato expression in neurons activated during the clozapine-labeling window.

After at least 4 weeks from 4-OHT administration, mice were challenged with clozapine, the receptor inhibitor cocktail, or olanzapine to assess drug-induced reactivation of the labeled ensemble. Mice were transcardially perfused 2 h after drug challenge, and brains were collected for c-Fos immunostaining.

For behavioral experiments requiring Cre-dependent viral expression, pAAV-EFS-FLEX-PSAM4-GlyR-HA-P2A-jGCaMP8f-WPRE was stereotaxically injected bilaterally into the medial prefrontal cortex (mPFC) of TRAP2 mice. The virus was diluted from 1.3 × 10^14 to 1.0 × 10^14 GC/ml before injection. A total volume of 500 nl virus was injected per side at the following coordinates: AP, +1.98 mm relative to bregma; ML, ±0.30 mm relative to bregma; and DV, −1.80 mm from the dura. Two weeks after viral injection, mice received intraperitoneal injections of clozapine (1 mg/kg) and 4-OHT (50 mg/kg) in the home cage to induce Cre recombination in clozapine-activated mPFC neurons. After at least 4 weeks from 4-OHT administration to allow stable transgene expression, mice were subjected to behavioral experiments. For chemogenetic inhibition experiments, uPSEM817 tartrate was administered intraperitoneally 30 min before behavioral testing.

### Histology

Mice were anesthetized with isoflurane, perfused with 4% paraformaldehyde, and decapitated. Brains were coronally sectioned at 40 μm using a cryostat. GFP was immunostained with anti-GFP antibody (1:1000, Santa Cruz, SC-9996). c-Fos was detected using an Alexa Fluor 647-conjugated anti-c-Fos antibody (rabbit monoclonal IgG, clone EPR21930-238, Abcam, ab300747). Secondary antibodies were goat anti-mouse IgG Alexa Fluor 488 (1:400, Jackson ImmunoResearch, #115-547-185). Nuclei were stained by DAPI. Sections were visualized under a Leica TCS SP8 confocal microscope.

### Quantification of TRAP2 and c-Fos overlap

TRAP2-labeled and drug-induced c-Fos-positive cells were quantified from maximum-intensity projection confocal images using a custom Python pipeline. DAPI, c-Fos, and tdTomato channels were used to identify nuclei, drug-induced activated cells, and TRAP2-labeled cells, respectively. Images were robustly normalized, Gaussian-smoothed, thresholded using Otsu’s method, and segmented by watershed-based object separation. Small objects were removed using channel-specific area thresholds of 50 pixels for DAPI, 70 pixels for c-Fos, and 100 pixels for tdTomato.

c-Fos/tdTomato overlap was determined from segmented object masks. A pair was classified as overlapping when the intersection area was ≥10 pixels and covered ≥30% of the c-Fos object and ≥20% of the tdTomato object. No mask dilation was applied before overlap detection. For each image series, DAPI-positive nuclei, c-Fos-positive cells, tdTomato-positive cells, and c-Fos/tdTomato overlap pairs were counted. Overlap was summarized as c-Fos-overlap fraction, tdTomato-overlap fraction, and overlap pairs normalized to DAPI-positive nuclei.

Cell segmentation and overlap quantification were performed using an automated pipeline with identical parameters across experimental groups. Treatment labels were not used during segmentation or overlap classification. Quality-control overlay images were generated to inspect segmentation and overlap detection.

### Behavioral task (probabilistic reversal-learning bandit task) and training procedure

#### Apparatus and task control

Behavioral experiments were conducted in custom-built operant chambers equipped with three nose-poke ports (center, left, right), each fitted with an infrared sensor and an LED cue. Water reward delivery was controlled by solenoid valves. Task control and data acquisition were implemented using the pyControl framework running on an Open Ephys–compatible behavioral control system (*39*).

All task logic was implemented as finite state machines written in Python (pyControl), with separate task scripts corresponding to each training stage (stages 1–5). Trial events, choices, reward outcomes, and block structure variables were logged on a trial-by-trial basis for subsequent analysis.

#### Water access and motivation strategy

To motivate task engagement while minimizing physiological stress, mice were not water restricted. Instead, mice had ad libitum access to citric acid–supplemented drinking water (2% w/v citric acid) in their home cages, following established protocols for maintaining stable motivation without dehydration (*40*). During behavioral sessions, plain water was used as the reward.

Body weight and general health were monitored throughout training and experimentation to ensure animals remained within normal physiological ranges.

#### Task structure: probabilistic reversal-learning bandit task

Mice were trained on a two-armed bandit task (2ABT) implemented in pyControl. Each session was controlled by a finite-state task program that registered left, center, and right port entries and delivered water rewards through left or right solenoid valves. Trials were composed of an initiation epoch, a choice epoch, an outcome epoch, and an inter-trial interval (ITI). During task stages that required trial initiation, the center port LED was illuminated at the beginning of each trial; a center-poke response extinguished the center LED and advanced the task to the choice epoch. During the choice epoch, the left and right port LEDs were illuminated, and the first left-or right-port entry was recorded as the animal’s choice. Reward delivery was determined immediately after the choice according to the current reward probabilities assigned to the chosen side, after which the task entered a 1-s ITI before the next trial.

Training proceeded through five task stages that gradually introduced the full probabilistic reversal structure. In stage 1, mice learned that water could be obtained from the two side ports without a center-poke initiation requirement. Both side ports were rewarded with probability 1.0, reward valves were opened for 100 ms, and sessions lasted 1 h. In stage 2, mice learned the complete trial sequence: each trial began with center-poke initiation, followed by a left/right choice, and both choices were rewarded with probability 1.0. Reward delivery lasted 100 ms and the ITI was 1 s. In stage 3, mice were introduced to a deterministic block structure. One side was designated as the high-value (good) side and the other as the low-value (bad) side; the good side was rewarded with probability 1.0 and the bad side with probability 0. After more than 30 successful choices of the good side, the rewarded side reversed after a one-trial delay. In stage 4, mice performed the same block-reversal structure, but the high-value side was rewarded probabilistically (0.95) while the low-value side remained unrewarded. In stage 5, the final probabilistic reversal task, sessions lasted 40 min, reward valves were opened for 50 ms, and the high- and low-value sides were rewarded with probabilities 0.80 and 0.20, respectively. After more than 30 choices of the current high-value side, a reversal was scheduled to occur after a randomly sampled delay of 1-10 trials. The first high-value side in each session was randomized between left and right. For each completed trial, the task program saved the trial number, cumulative rewards, block number, current high-value side, chosen side, reward outcome, and an exponential moving average of correct choices.

During testing, mice received vehicle, MK801, clozapine, or the inhibitor cocktail 30 min before the task according to a within-subject or between-subject design. In within-subject experiments, the order of drug administration was counterbalanced across animals.

### Reinforcement learning model and parameter estimation

Choice behavior was analyzed using a Q-learning framework (*15*) with asymmetric learning rates for positive and negative prediction errors (*41*), extended to explicitly model stochastic choice variability (“decision noise”) (*42*). Behavioral modeling was performed at the level of individual mice and drug conditions.

#### Q-value update rule

For each trial *t*, the action value *Q*_*t*_(*a*)for the chosen action *a* ∈ {left,right}was updated according to:

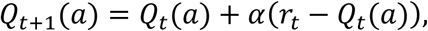

where *r*_*t*_ ∈ {0,1} denotes reward outcome. The learning rate *α* depended on the sign of the prediction error:

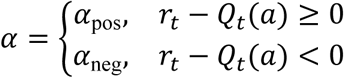

Separate learning rates were used to allow differential updating following rewarded and unrewarded outcomes, a feature commonly observed in rodent and human decision-making tasks. Initial Q-values were set to *Q*_0_(left) = *Q*_0_(right) = 0.5, reflecting unbiased initial expectations. Results were robust to alternative initializations (data not shown).

#### Action selection model with decision noise

Action selection followed an ε-softmax policy that combines value-based choice with random exploration (*43*):

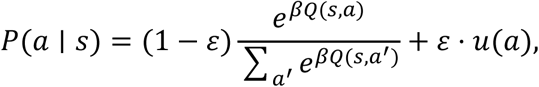

where *β* is the inverse temperature controlling choice determinism, *ε* represents decision noise, and *u*(*a*)is a uniform distribution over actions.

The parameter *ε*captures stochastic choice behavior that cannot be explained by value differences alone and was interpreted as a computational measure of decision-making noise. This formulation allows dissociation between value-guided choice (controlled by *β*) and value-independent randomness (controlled by *ε*).

For numerical stability, softmax probabilities were computed after subtracting the maximum Q-value at each trial.

#### Likelihood function

Model parameters were estimated by maximizing the log-likelihood of observed choices. For a sequence of actions {*a*_*t*_}and rewards {*r*_*t*_}, the negative log-likelihood was defined as:

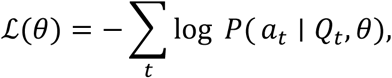

where *θ* = {*α*_pos_, *α*_neg_, *β*, *ε*}. A small constant (10^−12^) was added to action probabilities to avoid numerical issues arising from log (0).

#### Parameter optimization

Model parameters were estimated separately for each mouse and drug condition by minimizing the negative log-likelihood using the L-BFGS-B algorithm implemented in SciPy (scipy.optimize.minimize). Parameter bounds were imposed to ensure physiological plausibility and numerical stability:

- *α*_pos_, *α*_neg_ ∈ [0,1]
- *β* ∈ [0, *β*_max_], where *β*_max_ = 20
- *ε* ∈ [0, *ε*_max_], where *ε*_max_ = 0.6

Initial parameter values were set to *α*_pos_ = 0.3, *α*_neg_ = 0.2, *β* = 5.0, *ε* = 0.05.

#### Parameter recovery

Parameter recovery was performed to evaluate whether the model-fitting procedure could recover known generative values under the stage-5 task structure (*44*). The stage-5 reward probabilities and reversal rules were read directly from the pyControl task source, and synthetic sessions were generated using the same Q-learning-with-noise equations used for fitting. Unless changed at runtime, simulated agents used *α*_+_ = 0.3 and *α*_−_ = 0.2 Recovery simulations sampled a grid of three *ε* values (0.05, 0.3, and 0.55) and three *β* values (2, 8, and 16). For each grid point, multiple 300-trial sessions were simulated. During each simulated trial, the agent selected left or right according to the epsilon-softmax policy, reward was sampled from the stage-5 high- or low-value reward probability, and the chosen action value was updated according to the asymmetric learning rule. The stage-5 reversal rule was then applied: choices of the current high-value side incremented a success counter, and once the counter exceeded 30, the high-value side was reversed after a uniformly sampled 1-10 trial delay.

The synthetic trial tables were then fit with the same Q-learning-with-noise model used for empirical data. Recovery performance was summarized by comparing true and recovered values for *β* and *ε*, calculating recovery errors for each simulated condition, and generating vector PDF plots of true-versus-recovered parameters and recovery-error heatmaps. Successful recovery was interpreted as recovered estimates tracking the generative values across the parameter grid.

#### Posterior predictive validation and refitting of regenerated behavior

To assess whether fitted parameters could reproduce the observed behavioral structure, posterior predictive validation was performed using the fitted Q-learning-with-noise parameters from each mouse and condition. For each empirical mouse-by-condition fit, synthetic behavior was regenerated from the estimated *α*_+_, *α*_−_, *β*, and *ε* parameters. Whenever available, the empirical trial template was used to preserve the number of trials and the observed sequence of high-value sides; otherwise, a fallback stage-5 schedule was generated from the task rule. On each simulated trial, the fitted epsilon-softmax policy generated a choice, reward was sampled using the inferred or task-defined high- and low-value reward probabilities, and action values were updated with the fitted learning rates.

The regenerated behavioral datasets were re-analyzed. Each regenerated dataset was refit using the same Q-learning-with-noise model. Recovered parameters from the regenerated datasets were averaged across simulations and compared with the original empirical parameter estimates for the same source mouse and condition. This refit-after-regeneration procedure tested whether the fitted model was self-consistent: parameters estimated from real behavior should generate synthetic behavior that, when refit, yields similar parameter values and behavioral summary statistics.

### Fiber photometry recording

Fiber photometry recordings were performed using a custom-built pyPhotometry-based system (*45*). jGCaMP8f fluorescence was recorded in a two-excitation/one-emission configuration using time-division multiplexed illumination. A 405-nm LED (M405FP1, Thorlabs) was used as an isosbestic control channel, and a 470-nm LED (M470F4, Thorlabs) was used for calcium-dependent excitation. The 405-nm and 470-nm excitation light paths were filtered using a 405-nm bandpass filter (FBH405-10, Thorlabs) and a 469-nm bandpass filter (MF469-35, Thorlabs), respectively, and combined with a 490-nm dichroic mirror (DMSP490R, Thorlabs). Excitation light was delivered to the implanted optical fiber through a fiber-optic patch cable.

Emitted fluorescence was collected through the same fiber, separated from excitation light by the dichroic mirror, passed through a 525-nm emission filter (MF525-39, Thorlabs), and detected with an avalanche photodetector (APD440A2, Thorlabs). Signals were acquired using pyPhotometry software at 100 Hz. Behavioral events were controlled by pyControl and synchronized with photometry recordings by transistor-transistor logic (TTL) pulses sent from pyControl to the pyPhotometry acquisition system. TTL timestamps were used to align fluorescence signals to task events, including trial initiation, choice, reward delivery, and other task-defined events.

For chemogenetic inhibition experiments, uPSEM817 tartrate was administered intraperitoneally 30 min before behavioral testing. Photometry recordings were performed during the two-alternative bandit task while mice were freely moving.

### Photometry analysis

Fiber photometry data were analyzed using custom Python scripts. For each session, pyPhotometry .ppd files were paired with the corresponding pyControl behavioral log files. When possible, files were paired by matching filename stems; otherwise, timestamps parsed from filenames were used to identify the closest behavioral log for each photometry recording. Sessions lacking either a photometry file, a behavioral log file, or valid synchronization pulses were excluded from analysis.

Photometry signals were extracted from the .ppd files. Unless otherwise stated, the calcium-dependent fluorescence channel and the isosbestic control channel were processed as analog_2 and analog_1, respectively. The control channel was used to correct for non-calcium-dependent fluorescence fluctuations, and fluorescence changes were normalized as ΔF/F. Signals were low-pass filtered at 10 Hz before event-aligned analysis. Photometry timestamps were generated from the sampling rate recorded in the .ppd metadata.

Behavioral and photometry clocks were synchronized using TTL pulses sent from pyControl to the pyPhotometry acquisition system. Photometry-side synchronization pulse indices were converted to photometry timestamps and aligned to pyControl-side rsync event timestamps using Rsync-based clock alignment. All behavioral event times were transformed into the photometry time base before extracting event-aligned fluorescence traces.

Behavioral events were reconstructed from pyControl logs. For each valid trial, the analysis identified center-port entry, side choice, selected side, outcome time, reward outcome, high- versus low-value choice identity, and previous-trial outcome history. Trials without a defined reward outcome were excluded from event-aligned analyses. Fluorescence traces were aligned to center-port entry. Event-aligned traces were extracted from −2 s to +3 s around each event and averaged into 100-ms time bins. Unless otherwise specified, the pre-event interval was used as the baseline. In the primary analysis, event-aligned traces were normalized as z scores relative to the baseline period.

Events were assigned to multiple behavioral grouping schemes, including all trials, choice side, chosen value, reward outcome, value-by-outcome combinations, side-by-value-by-outcome combinations, and previous-to-current outcome history. High- versus low-value choices were assigned preferentially from the logged high-value side information. When this information was unavailable, logged left and right reward probabilities were used. Fixed block-size labeling was used only as a fallback.

For group-level traces, event-aligned fluorescence traces were first averaged within each session for each condition, mouse, grouping type, grouping label, and time bin. Group-level time courses were then calculated by averaging session-level means. The SEM at each time bin was computed across session-level means. These traces were used for summary plots and exported as long-format tables for reproducibility.

Photometry response magnitude was quantified as the area under the curve (AUC) within predefined peri-event windows. Unless otherwise stated, AUC was calculated from −0.5 to 0.5 s relative to the alignment event by trapezoidal integration of the event-aligned fluorescence signal. Event-level AUC values were averaged to obtain one session-level mean AUC for each condition, mouse, grouping type, grouping label, and session. These session-level AUC values were used for summary plots and statistical analyses.

Statistical comparisons of photometry AUC were performed using session-level mean AUC values. For each grouping type and grouping label, condition effects were tested using one-way ANOVA when more than two groups were compared. Pairwise comparisons were performed using Welch’s t-test. Within-condition comparisons between predefined trial labels were also performed using Welch’s t-test when applicable.

### Sleep recording and analysis

8-12-weeks-old wild-type mouse strains (C57BL/6J) were procured from CLEA Japan. Mice were housed under controlled temperature and humidity conditions and maintained on a 12-hour light: 12-hour dark cycle. EEG and EMG electrode implantation, EEG and EMG signal acquisition and data analyses were performed as previously described (*46*). Specifically, EEG electrodes were implanted at frontal (AP +1.0 mm, ML+1.5 mm from bregma) and occipital (AP −3.2 mm, ML+1.5 mm) areas. At 6 weeks of age, male mice were implanted with EEG and EMG electrodes under isoflurane anesthesia. Sleep recording began from two weeks post-surgery. The drugs were intraperitoneally injected at zeitgeber time 12. EEG and EMG were recorded at 128 Hz. Sleep stages were automatically classified using FASTER (*47*) at 8-s epochs. EEG signals were subjected to fast Fourier transform analysis to yield a power spectrum with a 0.39-Hz resolution. These spectra were normalized across the frequency axis. The relative power change was determined by dividing the power spectrum density by the baseline value. The baseline value represents the mean power from 5 hours before the intraperitoneal injection. The relative delta power change was the mean of the relative power change from 1 to 4 Hz.

**Fig. S1.**
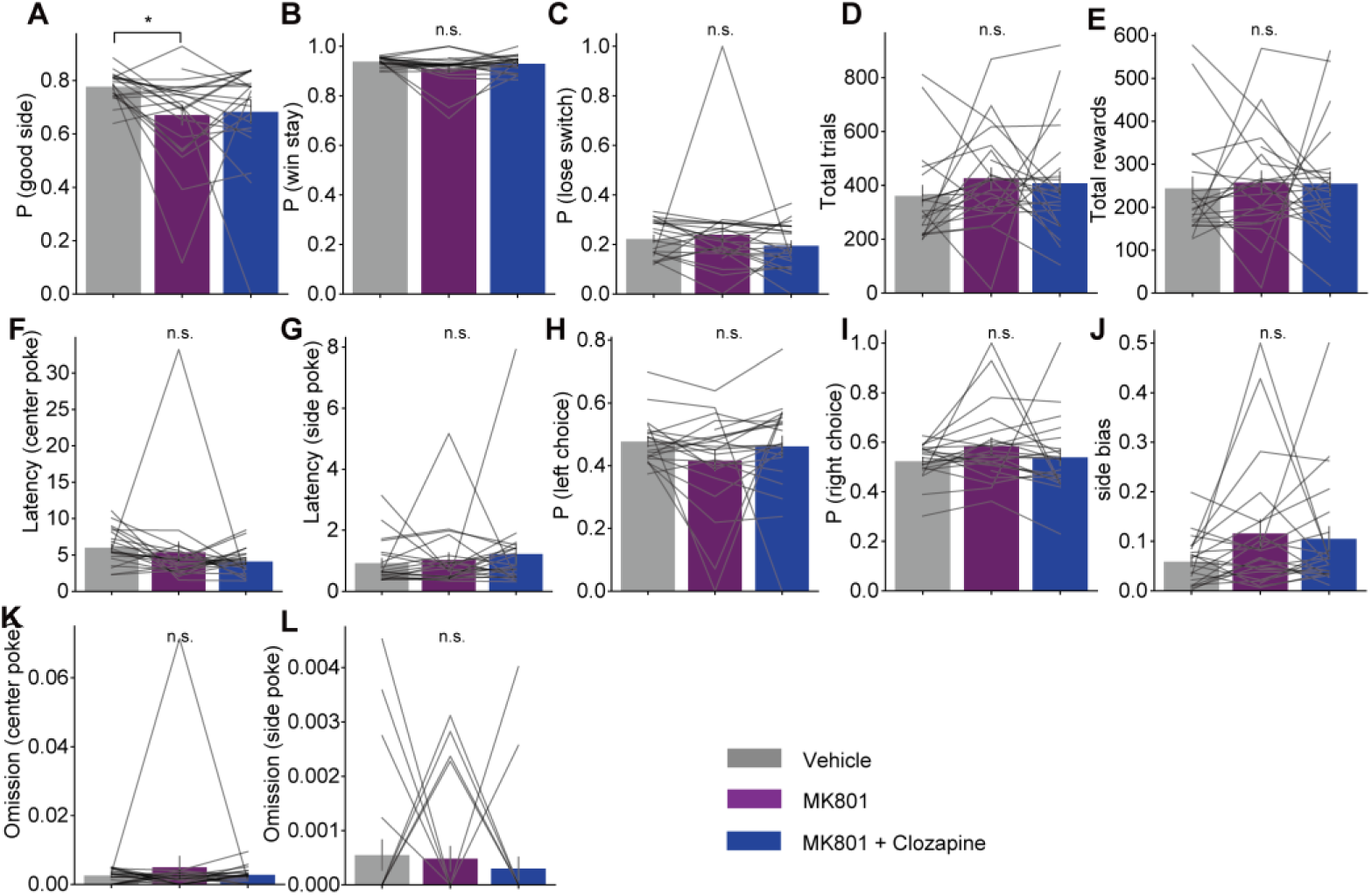
Additional behavioral metrics in the two-armed bandit task. **(A–L)** Secondary behavioral measures during the two-armed bandit task under saline, MK801, and MK801 plus clozapine conditions. Panels show the probability of choosing the currently better side (**A**), win-stay probability (**B**), lose-switch probability (**C**), total number of trials (**D**), total rewards obtained (**E**), latency from trial start to center poke (**F**), latency from center poke to choice (**G**), left-choice probability (**H**), right-choice probability (**I**), side bias (**J**), start-to-center omission rate (**K**), and center-to-choice omission rate (**L**). Bars indicate group means, and gray lines indicate paired individual mice (n = 21). Statistical comparisons were performed using one-way repeated-measures ANOVA followed by post hoc paired t tests with Holm correction. *P < 0.05; n.s., not significant.

**Fig. S2.**
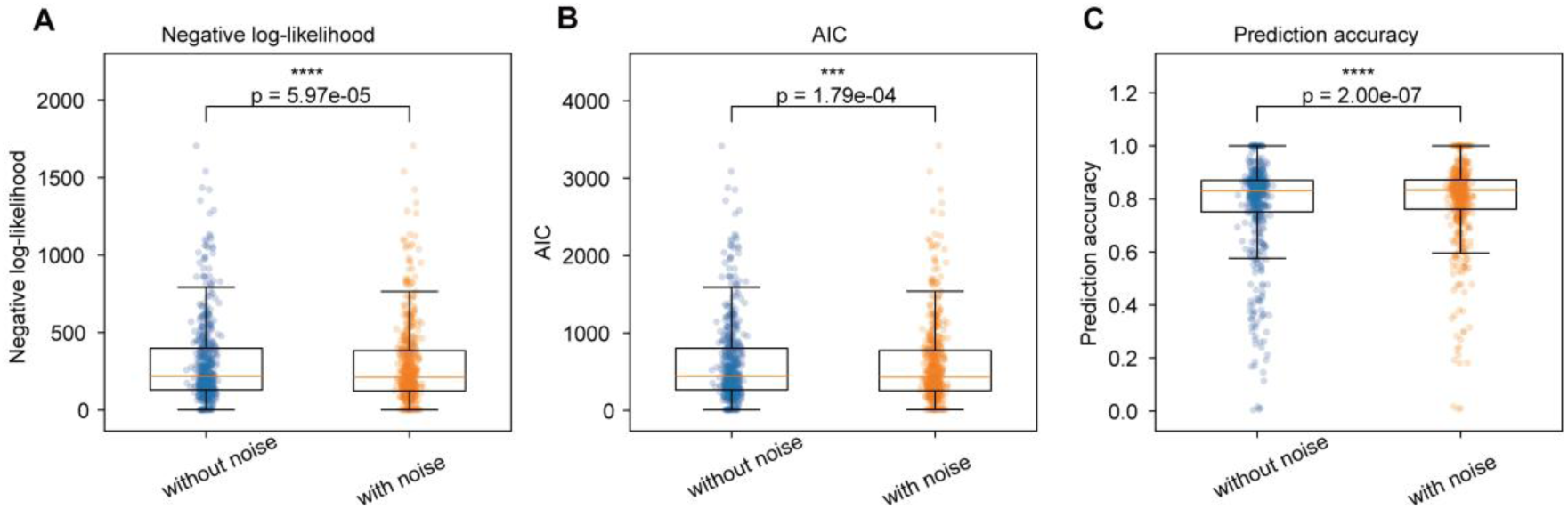
Q-learning model comparison supports inclusion of a value-independent noise parameter. **(A–C)** Comparison of Q-learning models fitted with or without a value-independent choice noise parameter, ε. Model performance was evaluated using negative log-likelihood (**A**), Akaike information criterion, AIC (**B**), and prediction accuracy (**C**). The model including ε showed significantly lower negative log-likelihood and AIC, and significantly higher prediction accuracy than the model without ε. Box plots show the distribution of fitted sessions, with individual data points overlaid (n = 519 sessions). Statistical comparisons were performed using two-sided paired t tests. ***P < 0.001; ****P < 0.0001.

**Fig. S3.**
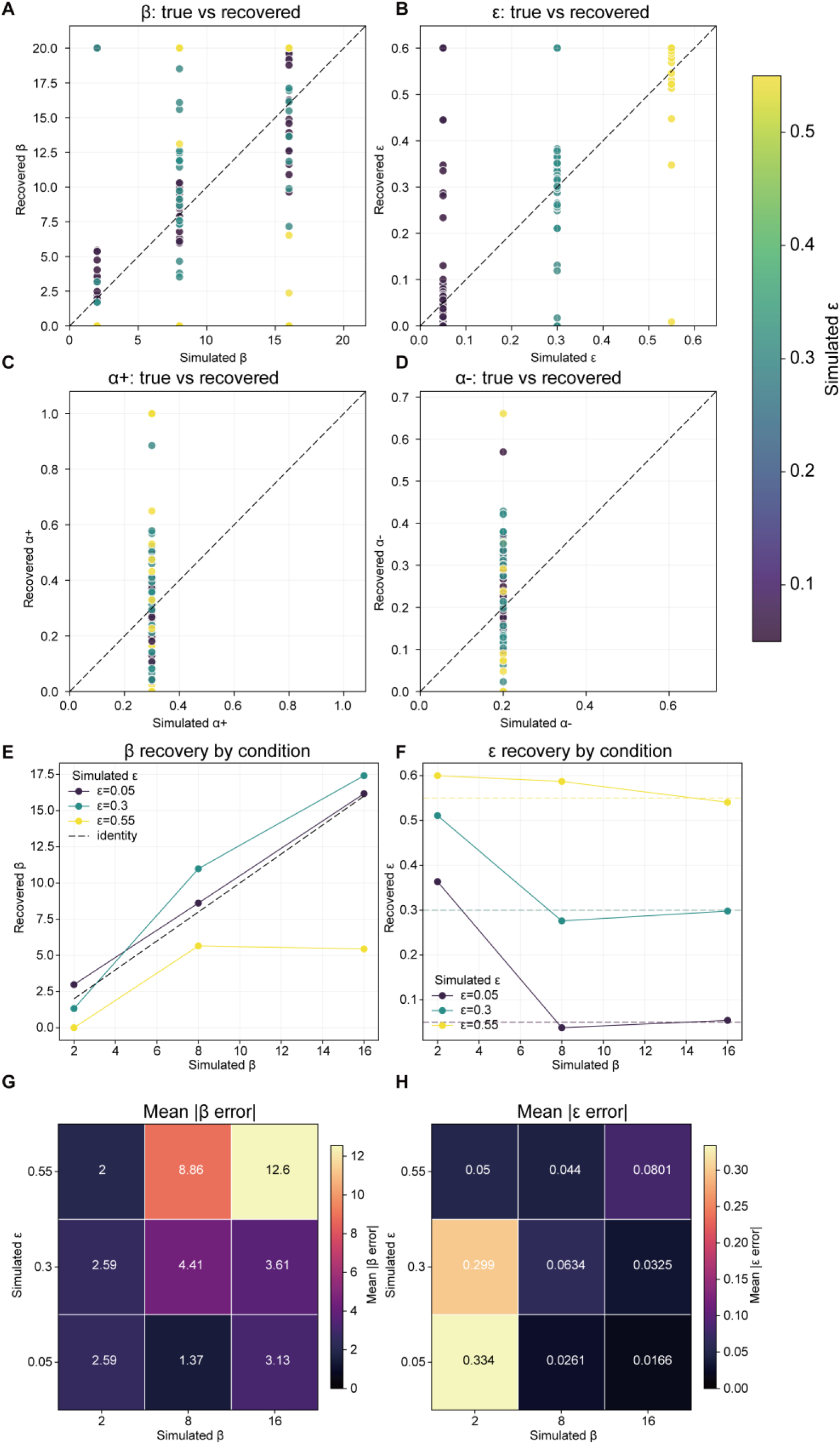
Parameter recovery of the Q-learning model with value-independent choice noise. **(A–D)** Parameter recovery analysis for the Q-learning model including a value-independent choice noise parameter, ε. Simulated behavioral datasets were generated across combinations of inverse temperature β and noise ε and then refitted with the same model. Recovered β (**A**), ε (**B**), positive learning rate α+(**C**), and negative learning rate α-(**D**) are plotted against the simulated values. Dashed lines indicate the identity line. Points are colored according to the simulated ε value. **(E, F)** Mean recovered β (**E**) and ε (**F**) across simulated β values, separately plotted for each simulated ε condition. Dashed horizontal or diagonal lines indicate the corresponding simulated parameter values. **(G, H)** Heatmaps showing the mean absolute recovery error for β (**G**) and ε (**H**) across simulated β and ε conditions.

**Fig. S4.**
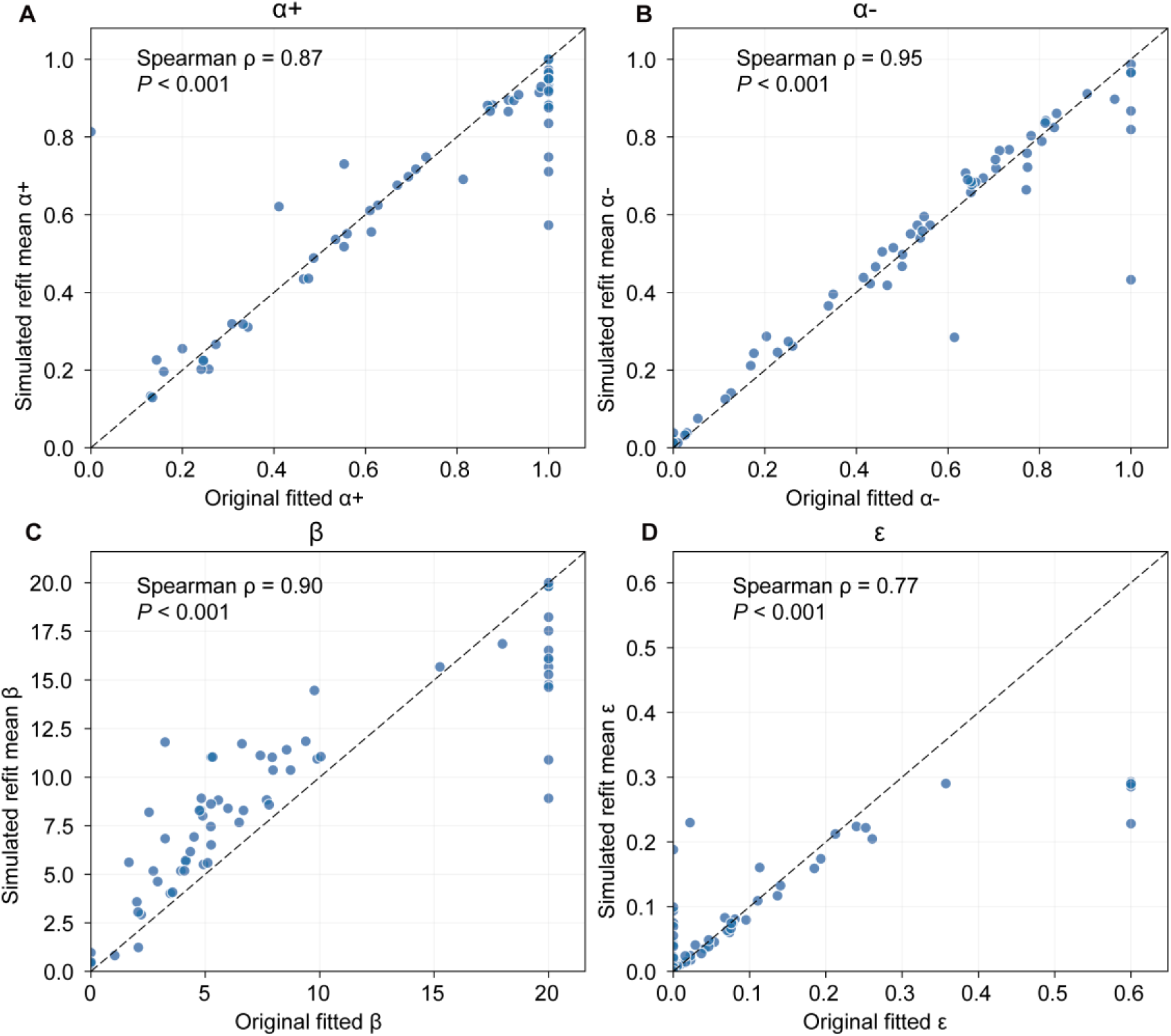
Simulation-based refitting supports recovery of empirically fitted Q-learning parameters. **(A–D)** Simulation-based parameter recovery using the empirically fitted Q-learning parameters. Behavioral datasets were simulated using the original fitted parameters from each session and then refitted with the same Q-learning model. The mean refitted parameters are plotted against the original fitted values for α+ (**A**), α− (**B**), β (**C**), and ε (**D**). Dashed lines indicate the identity line. The refitted α+ and α− closely tracked the original fitted values, indicating robust recovery of learning-rate parameters. β and ε also showed positive recovery trends, supporting their estimability within the empirical parameter range. However, recovery was less precise for high β and high ε values, indicating partial compression or underestimation in these parameter ranges. Spearman’s rank correlation was used to assess the association between ground-truth and refitted parameter values. P values are two-sided.

**Fig. S5.**
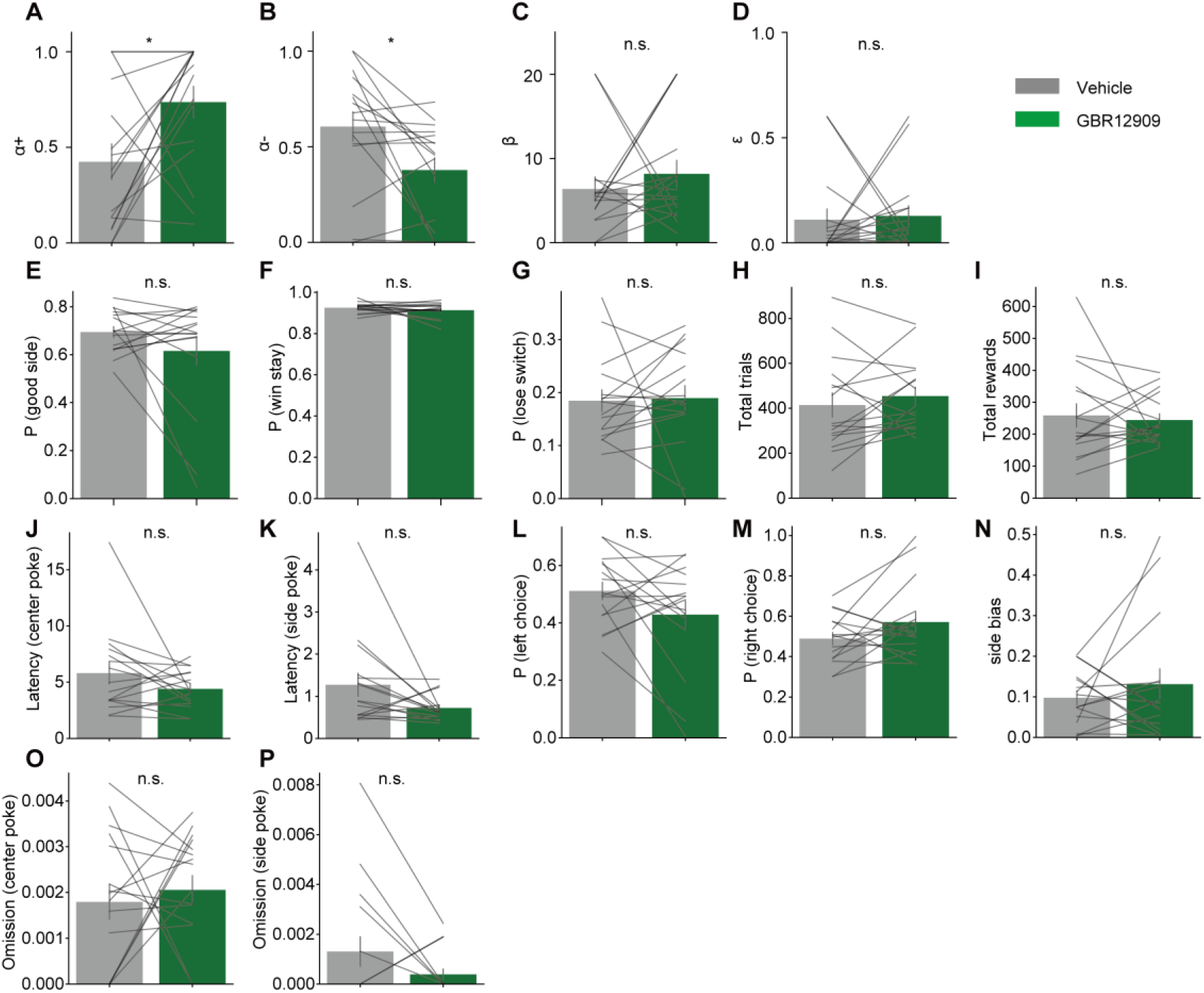
DAT inhibition selectively alters learning-rate parameters without producing broad changes in task performance. **(A–P)** Effects of the dopamine transporter inhibitor GBR12909 on fitted Q-learning parameters and behavioral measures in the two-armed bandit task. Panels show α+ (**A**), α− (**B**), β (**C**), ε (**D**), probability of choosing the better side (**E**), win-stay probability (**F**), lose-switch probability (**G**), total number of trials (**H**), total rewards obtained (**I**), latency from trial start to center poke (**J**), latency from center poke to choice (**K**), left-choice probability (**L**), right-choice probability (**M**), side bias (**N**), start-to-center omission rate (**O**), and center-to-choice omission rate (**P**). Gray and green bars indicate group means for vehicle and GBR12909 conditions, respectively, and thin gray lines indicate paired individual mice (n = 16). Among the fitted model parameters, GBR12909 significantly increased α+ and decreased α−, whereas β and ε were not significantly altered. No significant changes were detected in the other behavioral measures. Statistical comparisons were performed using two-sided paired t tests. *P < 0.05; n.s., not significant.

**Fig. S6.**
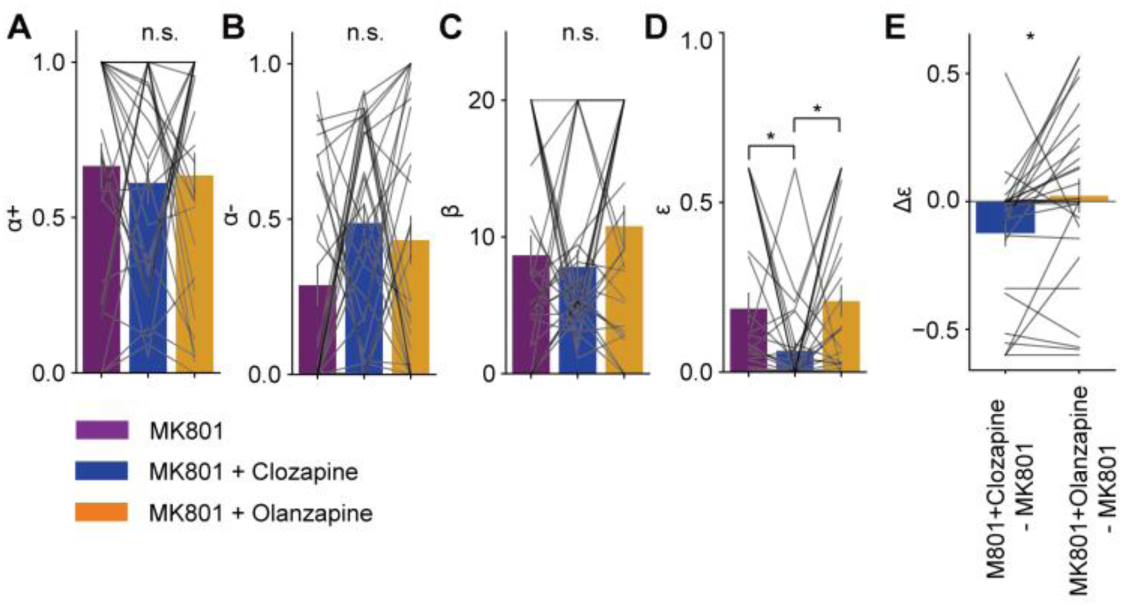
Olanzapine does not reproduce the clozapine-mediated rescue of MK801-induced decision-noise abnormalities. **(A–D)** Fitted Q-learning parameters in the two-armed bandit task under MK801, MK801 plus clozapine, and MK801 plus olanzapine conditions. Panels show α+ (**A**), α− (**B**), β (**C**), and ε (**D**). Bars indicate group means, and gray lines indicate paired individual mice (n = 28). **(E)** Within-subject comparison of changes in å relative to MK801. Clozapine reduced MK801-induced å, whereas olanzapine did not show a comparable reduction. The change in å was significantly different between MK801 plus clozapine and MK801 plus olanzapine conditions. Statistical comparisons for three-condition analyses were performed using one-way repeated-measures ANOVA followed by post hoc paired t tests with Holm correction. Two-condition comparisons were performed using two-sided paired t tests. *P < 0.05; n.s., not significant.

**Fig. S7.**
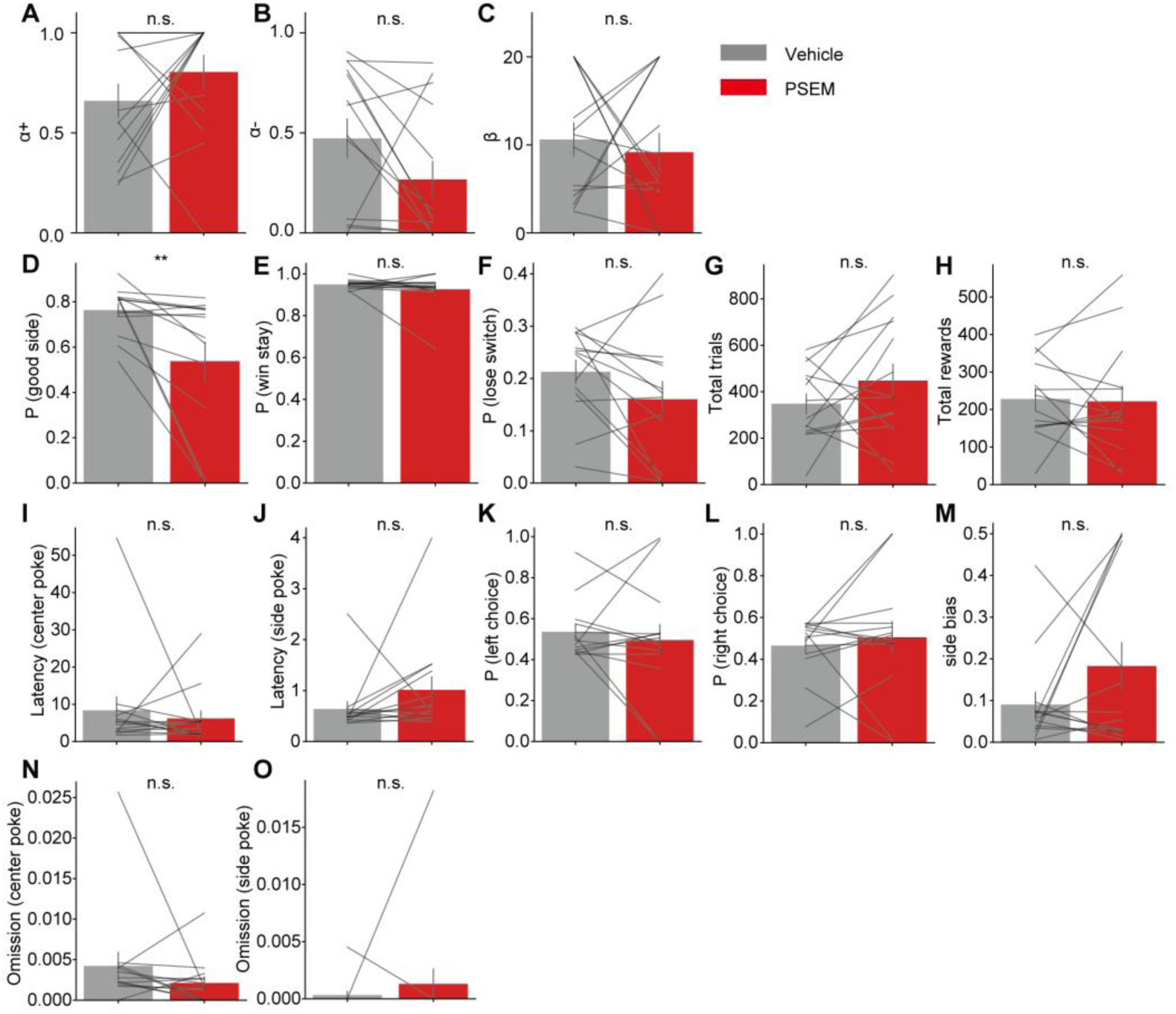
Additional behavioral measures during baseline inhibition of PFC clozapine-responsive cells. **(A–C)** Fitted Q-learning parameters during the two-armed bandit task under vehicle and PSEM conditions in mice with inhibition of PFC clozapine-responsive cells. Panels show α+ (**A**), α−(**B**), and β (**C**). **(D–O)** Secondary behavioral measures in the same task, including probability of choosing the better side (**D**), win-stay probability (**E**), lose-switch probability (**F**), total trials (**G**), total rewards (**H**), latency from trial start to center poke (**I**), latency from center poke to choice (**J**), left-choice probability (**K**), right-choice probability (**L**), side bias (**M**), start-to-center omission rate (**N**), and center-to-choice omission rate (**O**). Bars indicate group means, and gray lines indicate paired individual mice (n = 14). Statistical comparisons were performed using two-sided paired t tests. **P < 0.01; n.s., not significant.

**Fig. S8.**
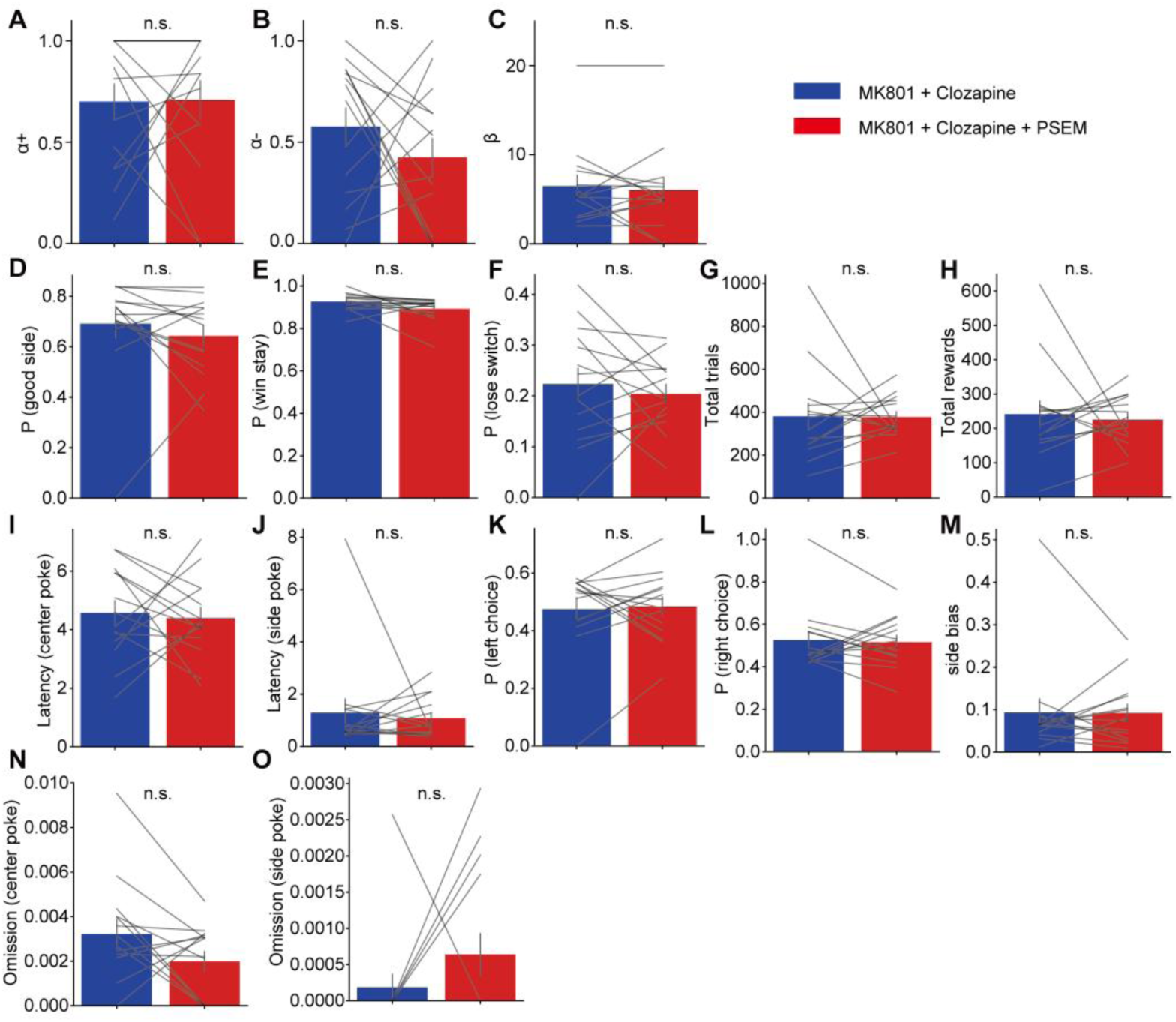
Additional behavioral measures during inhibition of PFC clozapine-responsive cells under MK801 plus clozapine treatment. **(A–C)** Fitted Q-learning parameters during the two-armed bandit task under MK801 plus clozapine and MK801 plus clozapine plus PSEM conditions in mice with inhibition of PFC clozapine-responsive cells. Panels show α+ (**A**), α− (**B**), and β (**C**). **(D–O)** Secondary behavioral measures in the same task, including probability of choosing the better side (**D**), win-stay probability (**E**), lose-switch probability (**F**), total trials (**G**), total rewards (**H**), latency from trial start to center poke (**I**), latency from center poke to choice (**J**), left-choice probability (**K**), right-choice probability (**L**), side bias (**M**), start-to-center omission rate (**N**), and center-to-choice omission rate (**O**). Bars indicate group means, and gray lines indicate paired individual mice (n = 14). Statistical comparisons were performed using two-sided paired t tests. n.s., not significant.

**Fig. S9.**
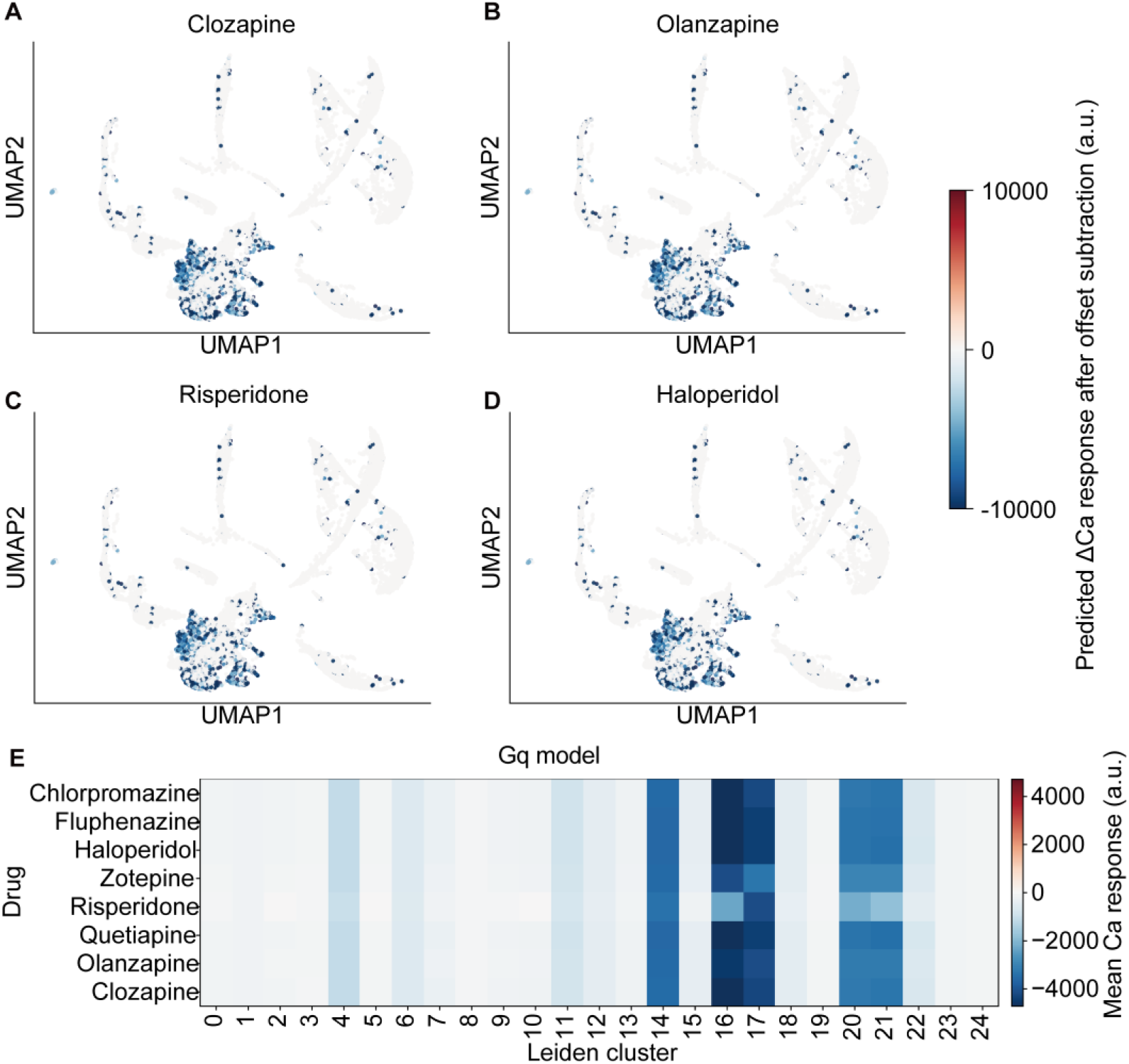
Gq-based Ca-response modeling shows limited drug selectivity across cell clusters. **(A–D)** UMAP visualization of predicted intracellular Ca-response changes (ΔCa) based on the Gq-signaling model for clozapine (**A**), olanzapine (**B**), risperidone (**C**), and haloperidol (**D**). Each point represents a single cell, colored by the predicted ΔCa response after offset subtraction. **(E)** Heatmap summarizing the mean predicted Ca response for each antipsychotic drug across Leiden clusters in the same dataset. Rows indicate drugs and columns indicate Leiden clusters. Color intensity represents the mean predicted Ca response (a.u.), with negative values indicating predicted suppression and positive values indicating predicted enhancement of Ca signaling.

**Fig. S10.**
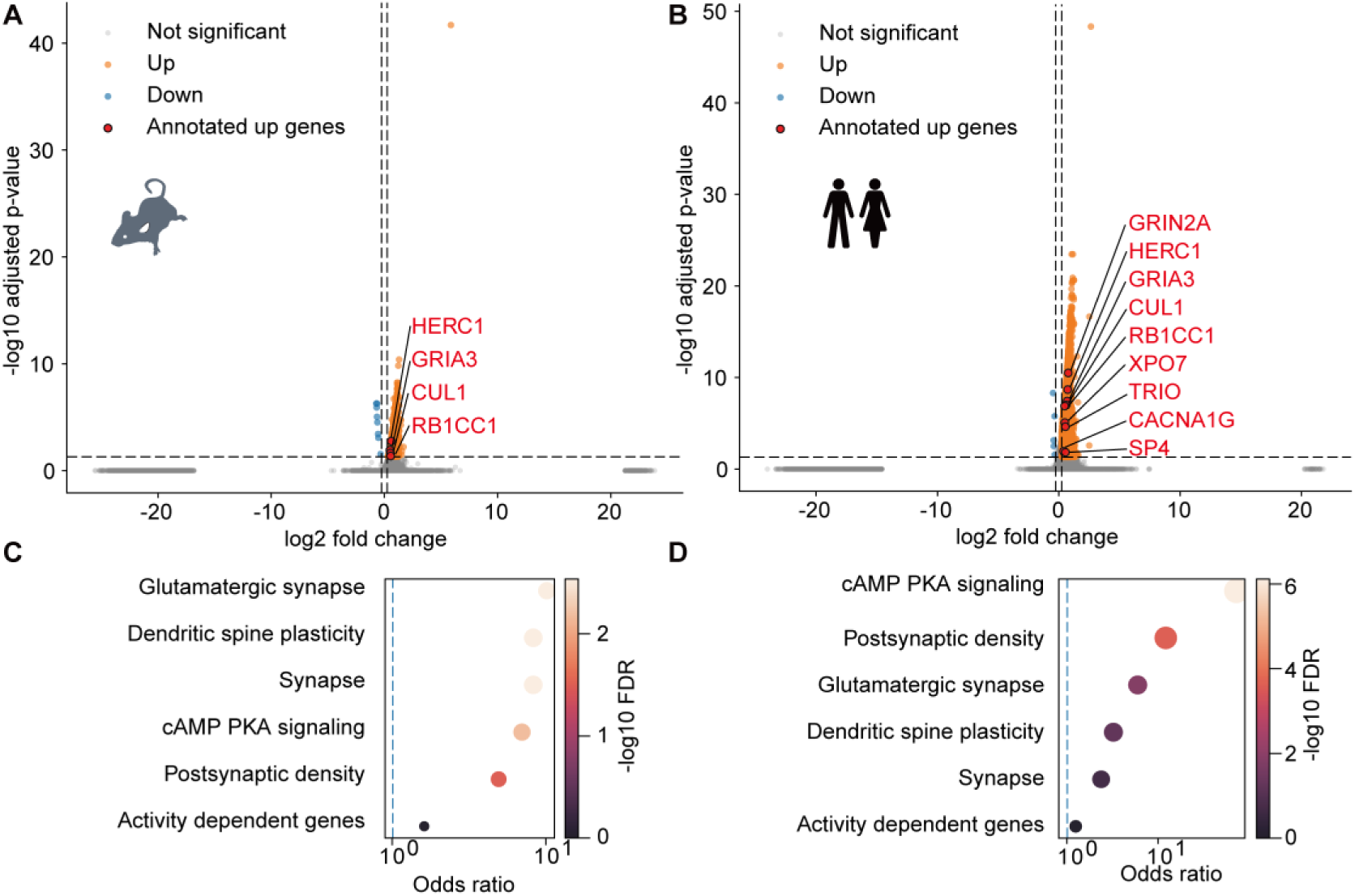
Excitatory-neuron-restricted DEG analysis preserves schizophrenia-risk and synaptic gene enrichment in clozapine-selective cells. **(A, B)** Volcano plots showing differentially expressed genes between predicted clozapine-selective and other excitatory neurons in mouse PFC (**A**) and human PFC (**B**). Orange and blue dots indicate significantly upregulated and downregulated genes, respectively. Red-labeled points indicate annotated SCHEMA schizophrenia risk genes among genes upregulated in clozapine-selective excitatory neurons. Dashed lines indicate the thresholds for adjusted P value and log2 fold change. **(C, D)** Curated gene-set enrichment analysis of genes upregulated in clozapine-selective excitatory neurons in mouse (**C**) and human (**D**). Gene sets related to schizophrenia risk, glutamatergic synapses, postsynaptic density, dendritic spine plasticity, synapses, cAMP/PKA signaling, and activity-dependent genes were tested. Dot position indicates the odds ratio, and color indicates −log10 FDR.

**Fig. S11.**
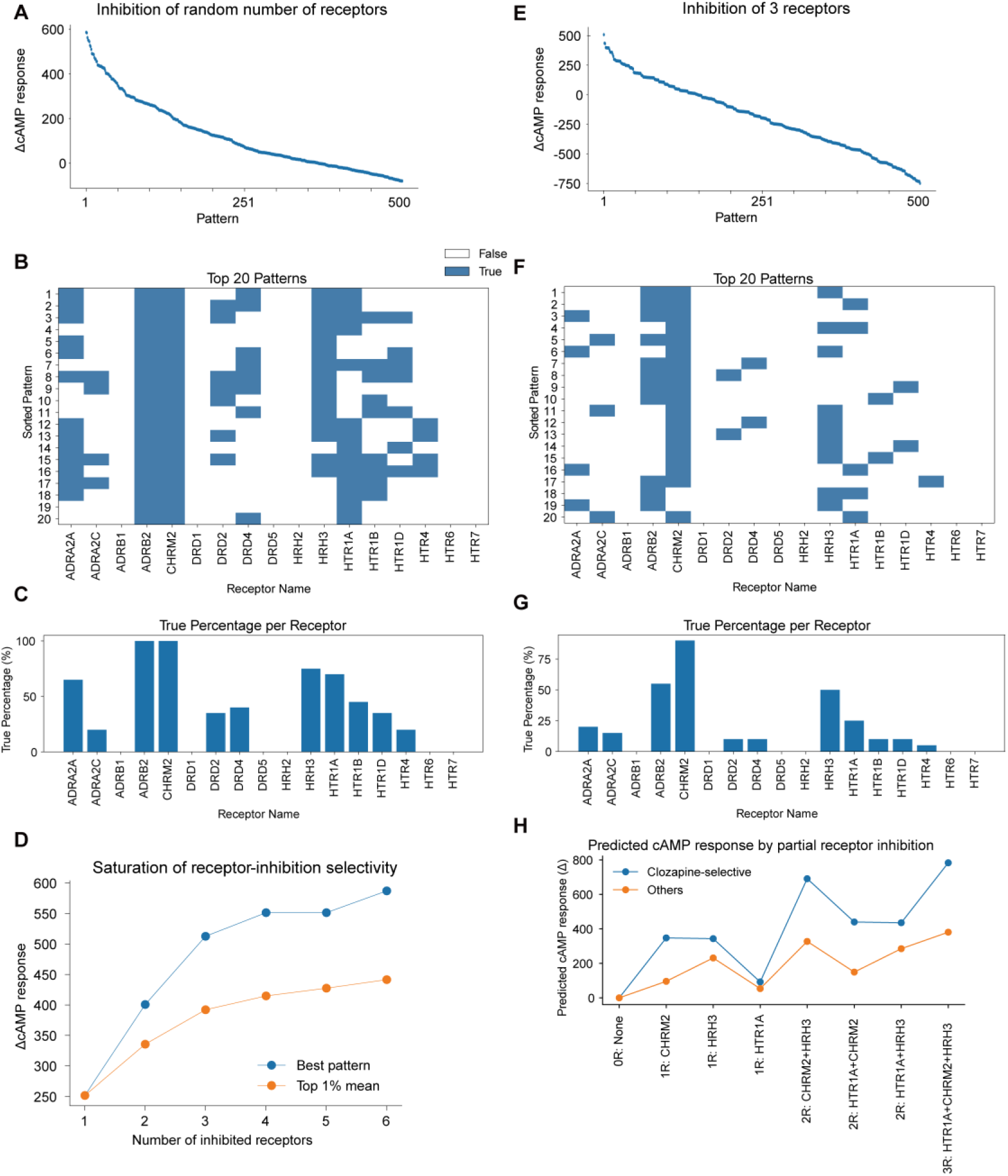
Rational design of receptor-inhibition patterns that selectively activate clozapine-selective cells. **(A–D)** Screening of receptor-inhibition patterns to identify combinations predicted to preferentially increase cAMP signaling in clozapine-selective cells relative to other cells. **(A)** Ranked distribution of predicted ÄcAMP selectivity scores obtained from patterns containing variable numbers of inhibited receptors. **(B)** Receptor composition of the top 20 patterns from this screen. Blue indicates receptors included in each inhibition pattern. **(C)** Frequency with which each receptor appeared among the top-ranked patterns. **(D)** Relationship between the number of inhibited receptors and predicted selectivity, shown as the best-performing pattern and the mean of the top 1% of patterns for each receptor-number condition. These analyses indicate a saturating gain in selectivity as the number of inhibited receptors increases. **(E–H)** Focused analysis of patterns containing exactly three inhibited receptors. **(E)** Ranked distribution of predicted ΔcAMP selectivity scores for all three-receptor inhibition patterns. **(F)** Receptor composition of the top 20 three-receptor patterns. **(G)** Frequency with which each receptor appeared among the top-ranked three-receptor patterns. **(H)** Stepwise comparison of predicted cAMP responses in clozapine-selective and other cells for partial and complete combinations of the final candidate receptors. Sequential addition of **CHRM2**, **HRH3**, and **HTR1A** increased the predicted response preferentially in clozapine-selective cells, supporting this triple inhibition pattern as a rationally selected cocktail.

**Fig. S12.**
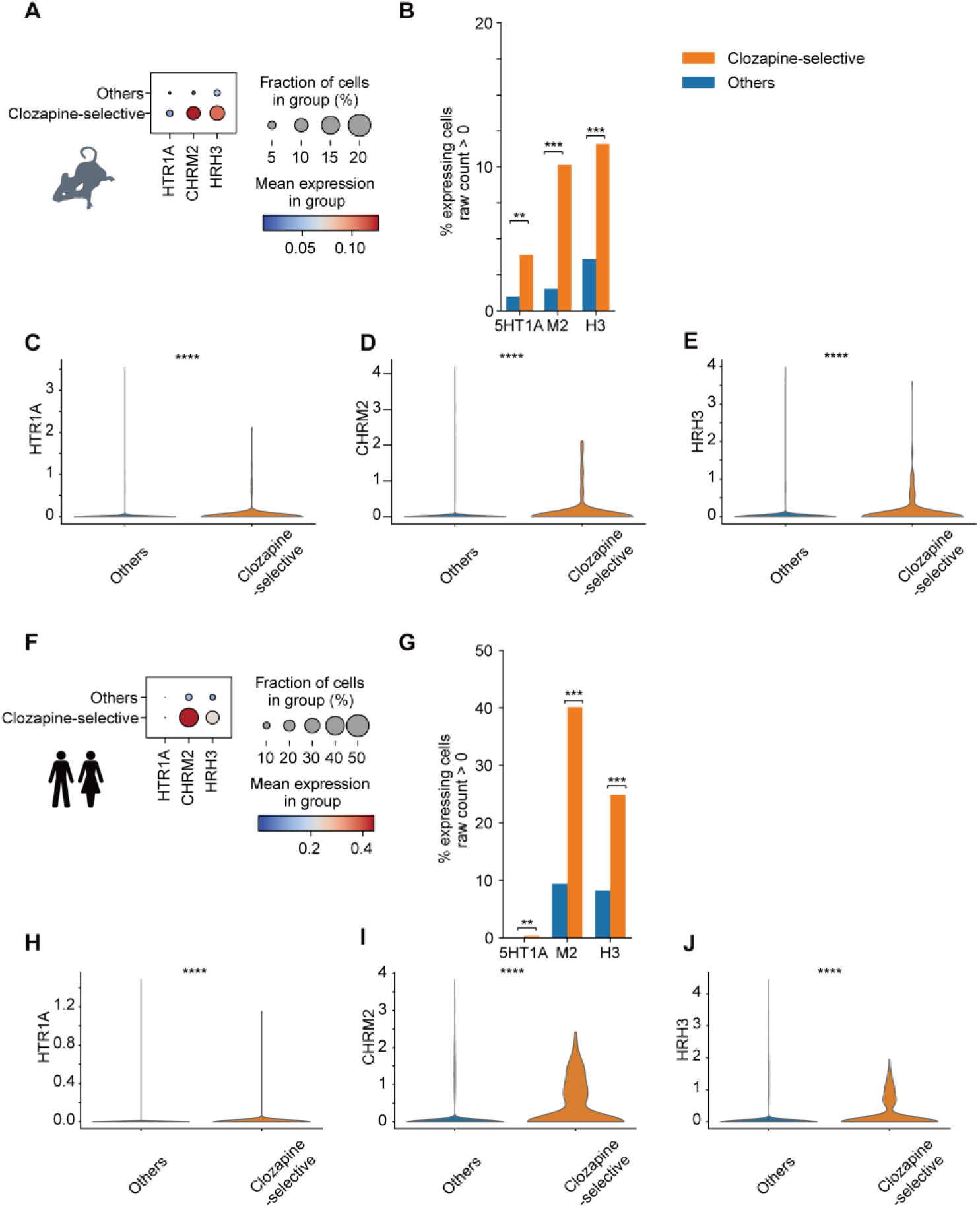
Candidate receptor expression in mouse and human clozapine-selective cells. **(A–E)** Expression analysis of the candidate receptors **HTR1A**, **CHRM2**, and **HRH3** in mouse PFC cells. **(A)** Dot plot showing receptor expression in predicted clozapine-selective cells and other cells. Dot size indicates the fraction of cells expressing each receptor, and color indicates mean expression level within each group. **(B)** Percentage of cells with detectable raw-count expression of each receptor in clozapine-selective cells and other cells. q values indicate multiple-comparison-adjusted significance. **(C–E)** Violin plots showing expression distributions of **HTR1A** (**C**), **CHRM2** (**D**), and **HRH3** (**E**) in clozapine-selective cells and other cells. **(F–J)** Corresponding receptor-expression analysis in human PFC cells. **(F)** Dot plot showing expression of **HTR1A**, **CHRM2**, and **HRH3** in predicted clozapine-selective cells and other cells. **(G)** Percentage of cells with detectable raw-count expression of each receptor in clozapine-selective cells and other cells. q values indicate multiple-comparison-adjusted significance. **(H–J)** Violin plots showing expression distributions of **HTR1A** (**H**), **CHRM2** (**I**), and **HRH3** (**J**) in clozapine-selective cells and other cells. Differences in expression levels were tested using two-sided Mann–Whitney U tests, and differences in the fraction of expressing cells were tested using Fisher’s exact tests. P values were adjusted for multiple comparisons using the Benjamini–Hochberg false discovery rate procedure. q values indicate FDR-adjusted P values. **q < 0.01; ***q < 0.001; ****q < 0.0001.

**Fig. S13.**
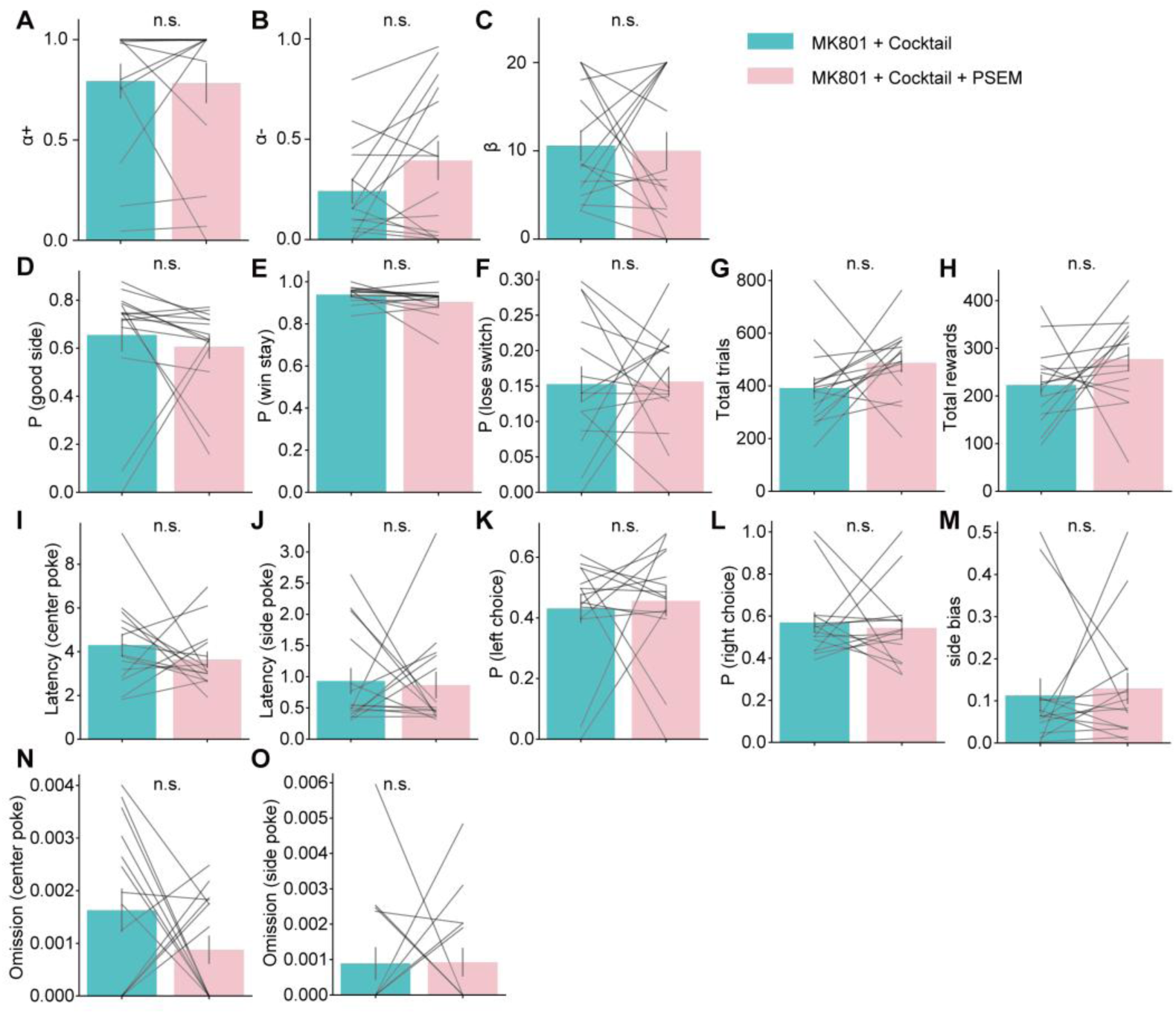
Additional behavioral measures during inhibition of PFC clozapine-responsive cells under MK801 plus cocktail treatment. **(A–C)** Fitted Q-learning parameters during the two-armed bandit task under MK801 plus cocktail and MK801 plus cocktail plus PSEM conditions in mice with inhibition of PFC clozapine-responsive cells. Panels show α+ (**A**), α− (**B**), and β (**C**). **(D–O)** Secondary behavioral measures in the same task, including probability of choosing the better side (**D**), win-stay probability (**E**), lose-switch probability (**F**), total trials (**G**), total rewards (**H**), latency from trial start to center poke (**I**), latency from center poke to choice (**J**), left-choice probability (**K**), right-choice probability (**L**), side bias (**M**), start-to-center omission rate (**N**), and center-to-choice omission rate (**O**). Bars indicate group means, and gray lines indicate paired individual mice (n = 15). Statistical comparisons were performed using two-sided paired t tests. n.s., not significant.

**Fig. S14.**
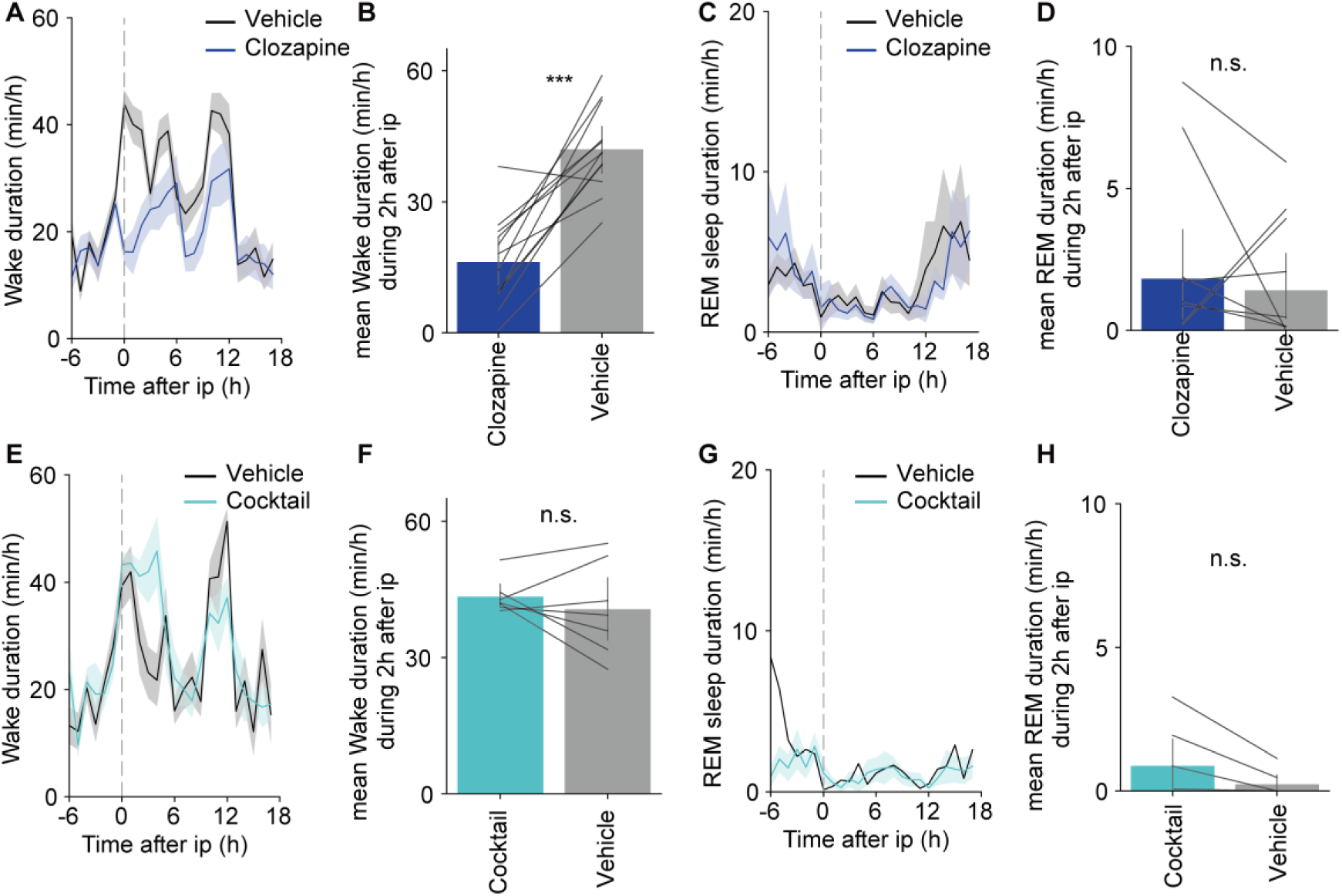
Effects of clozapine and the receptor-inhibition cocktail on wakefulness and REM sleep. **(A–D)** Effects of clozapine on wakefulness and REM sleep. Time-course plots show wake duration (**A**) and REM sleep duration (**C**) before and after intraperitoneal injection. Dashed vertical lines indicate the time of injection. Shaded areas indicate SEM. Summary plots show mean wake duration (**B**) and mean REM sleep duration (**D**) during the 2 h after injection. Clozapine significantly reduced wake duration compared with vehicle, whereas REM sleep duration was not significantly altered. n = 12 mice. **(E–H)** Effects of the receptor-inhibition cocktail on wakefulness and REM sleep. Time-course plots show wake duration (**E**) and REM sleep duration (**G**) before and after injection. Summary plots show mean wake duration (**F**) and mean REM sleep duration (**H**) during the 2 h after injection. n = 7 mice. Bars indicate group means, and gray lines indicate paired individual mice. Statistical comparisons were performed using two-sided Wilcoxon signed-rank tests. ***P < 0.001; n.s., not significant.

**Data S1. (separate file)**

Complete differential expression results for mouse predicted clozapine-selective cells related to Fig. 3K. The table includes gene-level differential expression statistics for predicted clozapine-selective cells compared with non-selective cells in the mouse prefrontal cortex.

**Data S2. (separate file)**

Complete differential expression results for human predicted clozapine-selective cells related to Fig. 3L. The table includes gene-level differential expression statistics for predicted clozapine-selective cells compared with non-selective cells in the human prefrontal cortex.

**Data S3. (separate file)**

Complete differential expression results for mouse excitatory predicted clozapine-selective cells related to fig. S10A. The table includes gene-level differential expression statistics restricted to excitatory cells in the mouse prefrontal cortex.

**Data S4. (separate file)**

Complete differential expression results for human excitatory predicted clozapine-selective cells related to fig. S10B. The table includes gene-level differential expression statistics restricted to excitatory cells in the human prefrontal cortex.

